# Modeling spatial evolution of multi-drug resistance under drug environmental gradients

**DOI:** 10.1101/2023.11.16.567447

**Authors:** Tomas Freire, Zhijian Hu, Kevin B. Wood, Erida Gjini

**Affiliations:** Center for Computational and Stochastic Mathematics, Instituto Superior Técnico, University of Lisbon, Lisbon, Portugal; Departments of Biophysics and Physics, University of Michigan, USA

**Keywords:** infectious diseases, spatial evolutionary dynamics, multi-drug antibiotic resistance, environmental heterogeneity, selection

## Abstract

Multi-drug combinations to treat bacterial populations are at the forefront of approaches for infection control and prevention of antibiotic resistance. Although the evolution of antibiotic resistance has been theoretically studied with mathematical population dynamics models, extensions to spatial dynamics remain rare in the literature, including in particular spatial evolution of multi-drug resistance. In this study, we propose a reaction-diffusion system that describes the multi-drug evolution of bacteria, based on a rescaling approach (Gjini and Wood, 2021). We show how the resistance to drugs in space, and the consequent adaptation of growth rate is governed by a Price equation with diffusion. The covariance terms in this equation integrate features of drug interactions and collateral resistances or sensitivities to the drugs. We study spatial versions of the model where the distribution of drugs is homogeneous across space, and where the drugs vary environmentally in a piecewise-constant, linear and nonlinear manner. Applying concepts from perturbation theory and reaction-diffusion equations, we propose an analytical characterization of *average mutant fitness* in the spatial system based on the principal eigenvalue of our linear problem. This enables an accurate translation from drug spatial gradients and mutant antibiotic susceptibility traits, to the relative advantage of each mutant across the environment. Such a mathematical understanding allows to predict the precise outcomes of selection over space, ultimately from the fundamental balance between growth and movement traits, and their diversity in a population.

## 1. Introduction

Bacterial resistance to antibiotics remains one of the biggest threats to public health. The emergence and selection of strains that are resistant to multiple antibiotics exacerbates the problem of resistance management and control (Laxminarayan et al., 2013; Cassini et al., 2019; Marston et al., 2016; Murray et al., 2022). Different strategies have been proposed to mitigate the problem of antibiotic resistance, including antibiotic cycling vs. mixing patterns (Beardmore et al., 2017; Brown and Nathwani, 2005; Bergstrom et al., 2004; Nichol et al., 2015; Batra et al., 2021), synergies with the host immune defenses (Gjini and Brito, 2016), maintenance of competition with sensitive strains (Hansen et al., 2017, 2020; Gatenby et al., 2009; West et al., 2020), and–crucially–multidrug evolutionary strategies (Baym et al., 2016). Although the molecular mechanisms underlying the rapid evolution of drug resistance are increasingly understood, it remains difficult to link this molecular and genetic information with multi-species population dynamics at different scales (MacLean et al., 2010; Holmes et al., 2016; Singer et al., 2007; Denk-Lobnig and Wood, 2023). In addition, while the majority of studies focus on bacteria populations in well-mixed environments (Lane et al., 1999; Kawecki et al., 2012; Hughes and Andersson, 2017), natural communities evolve on spatially extended habitats that display multiple biotic and abiotic gradients (Donaldson et al., 2016; Chikina and Vignjevic, 2021) and potentially yield complicated networks of interacting subpopulations (Hanski, 1998; Nicoletti et al., 2023).

Understanding the role of this spatial structure in the evolution of resistance is an ongoing challenge, despite the fact that environmental drivers of evolution–for example, the local concentration of antibiotic, or the density of susceptible hosts–are known to vary on multiple scales–across different body compartments, organs, or tissues, and on longer length scales, between hospitals and geographic regions.

The impact of spatial heterogeneity on ecological and evolutionary dynamics has been studied in a wide range of contexts–from the spread of COVID (Thomas et al., 2020) and HIV (Zulu et al., 2014; Feder et al., 2021) to conservation ecology (Silva et al., 2006; Hovick et al., 2015). Graph theory and dynamical systems theory offer a number of elegant approaches for studying multi-habitat models on networks (Allen et al., 2017; Marrec et al., 2021) or in particular limits (e.g. with a center manifold reduction) (Constable and McKane, 2014). Theory indicates that the way different subpopulations in a community are topologically “connected” can alter evolution (Lieberman et al., 2005; Marrec et al., 2021), and laboratory experiments in microbes are beginning to confirm some of these predictions (Kreger et al., 2023; Chakraborty et al., 2021). In parallel, a separate body of work has focused on spatial dynamics of microbial communities on agar plates, leading to an increasingly mature understanding of range expansions and cooperation in multi-species communities (Korolev et al., 2010; Korolev, 2013, 2015; Datta et al., 2013; Sharma and Wood, 2021; Martínez-Calvo et al., 2023) or in populations impacted by complex fluid dynamics Atis et al. (2019); Plummer et al. (2019).

In the specific context of drug resistance, spatial heterogeneity can manifest in multiple ways, from heterogeneity in drug concentrations to host heterogeneity in infectious disease models Brockhurst et al. (2004); Campos et al. (2008); Chabas et al. (2018). A number of theoretical and experimental studies have shown that spatial differences in drug concentration significantly impact the evolution of resistance (Galvin et al., 2013; Organization et al., 2022; Fu et al., 2015; Greulich et al., 2012; Hermsen et al., 2012; Kepler and Perelson, 1998; Moreno-Gamez et al., 2015; Zhang et al., 2011; Hermsen and Hwa, 2010; Baym et al., 2016; De Jong and Wood, 2018a). Theory suggests that the presence of spatial gradients of drug tends to accelerate resistance evolution (Hermsen and Hwa, 2010; Hermsen et al., 2012; Kepler and Perelson, 1998; Moreno-Gamez et al., 2015; Fu et al., 2015), though it can be slowed down by tuning the drug profiles (De Jong and Wood, 2018b) or in cases where the fitness landscape is non-monotonic Greulich et al. (2012).

The connection between spatial heterogeneity and the evolution of resistance is particularly murky when multiple drugs are involved, making predictions difficult, both at a within-host level in a clinical setting, as well as at the higher population or ecological levels (Goossens et al., 2005; Asaduzzaman et al., 2022). Even in the absence of spatial structure, multi-drug therapies are a subject of intense interest. Antibiotics interact when the combined effect of the drugs is greater than (synergy) or less than (antagonism) expected based on the effects of the drugs alone (Loewe, 1953; Greco et al., 1995), and these interactions can accelerate, reduce, or even reverse the evolution of resistance (Chait et al., 2007; Michel et al., 2008; Hegreness et al., 2008; Pena-Miller et al., 2013; Dean et al., 2020; Gjini and Wood, 2021). In addition to these interactions, which occur when drugs are used simultaneously, resistance to different drugs is linked through collateral effects–where resistance to one drug is associated with modulated resistance to other drugs. Collateral effects have been recently shown to significantly modulate resistance evolution (Barbosa et al., 2018; Rodriguez de Evgrafov et al., 2015; Munck et al., 2014; Maltas and Wood, 2019; Maltas et al., 2020; Roemhild et al., 2020; Ardell and Kryazhimskiy, 2021).

Despite substantial progress in understanding spatial heterogeneity, drug interactions, and collateral effects separately, it remains unclear how these three components combine to impact the evolution of multi-drug antibiotic resistance. To address this gap, we study a general mathematical model describing a continuous environment with spatially-varying antibiotic concentrations, in which bacteria move, grow and are selected following deterministic dynamics. Our aim is to build an integrative framework for drug interactions and collateral effects in spatially-extended multi-drug environments and show how specific drug gradients can shape evolutionary outcomes.

The outline of the paper is as follows. In Section 2 we present the spatial model, extending the model–first introduced in (Gjini and Wood, 2021) for multi-drug resistance evolution (summarized in BOX 1). In Section 3, we present an analytical quantity for predicting outcomes of selection in multi-drug spatial environments, and describe key cases of multi-drug resistance evolution linking simulations with theoretical predictions. We conclude with a discussion of our study’s limitations and potential future extensions.

### BOX 1.

Modeling framework for multidrug resistance

#### Drug resistance as a rescaling of effective drug concentration

To link a cell’s level of antibiotic resistance with its fitness in a given multidrug environment, we assume that drug-resistant mutants exhibit phenotypes identical to those of the ancestral (“wild type”) cells but at rescaled effective drug concentration (Chait et al., 2007; Gjini and Wood, 2021). The phenotypic response (e.g. growth rate) of drug-resistant mutants corresponds to a rescaling of the growth rate function *G*(*x, y*), of the ancestral population at concentrations *x* and *y* of the two drugs. At such concentrations, the per-capita growth rate (*g*_*i*_) of mutant *i* is given by

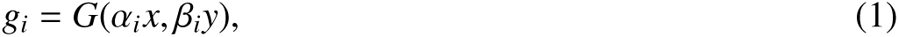

where *α*_*i*_ and *β*_*i*_ are rescaling parameters that reflect an effective change in drug concentration and, therefore, in that mutant’s subpopulation growth rate.

##### Mutant traits

In a 2-drug environment, each mutant is characterized by a pair of scaling parameters,

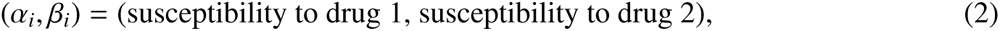

which one might think of as a type of coarse-grained genotype. They can be measured experimentally, for example, as the ratios of the mutant MICs for two different antibiotics relative to those of the wild-type.

##### Mean trait evolution and population adaptation

Considering all growth rates *g*_*i*_ of existing mutants (e.g. *i* = 1, ..*M*), the population dynamics of scaling parameters follows naturally as a dynamic weighted average over all sub-populations. The mean resistance traits to drugs 1 and 2, (averaged over all mutants)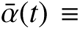 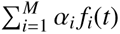, and 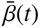 evolve as:

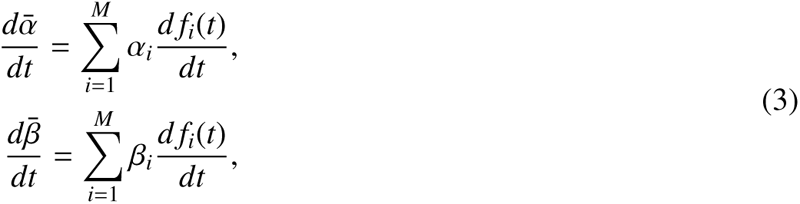

where *f*_*i*_(*t*) is the frequency of mutant *i* at time *t* in the population. Assuming exponential growth (*dn*_*i*_/*dt* = *g*_*i*_*n*_*i*_, with *n*_*i*_ the abundance of mutant *i* and *g*_*i*_ given by Equation 1), the frequency *f*_*i*_(*t*) changes as:

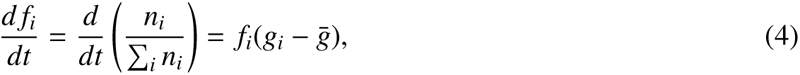

where 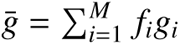 is the (time-dependent) mean value of *g*_*i*_ across all *M* subpopulations (mutants).

##### The Price Equation for mean trait evolution

Combining equations, we arrive at:

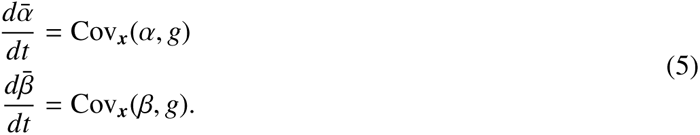

where 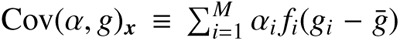 is the covariance between the scaling parameters α_*i*_ and the corresponding mutant growth rates *g*_*i*_, and similarly for Cov(β, *g*)_***x***_. The subscript ***x*** refers to the fact that the growth rates *g*_*i*_ and 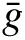 depend on the external (true) drug concentration ***x*** ≡ (*x, y*).

## 2. Methods

### 2.1. The model extended to space and spatial gradients of 2 drugs

We consider the case of a population of bacteria growing and diffusing in space (1-d) as a finite set of *M* subpopulations (mutants/strains), where each mutant has a potentially different level of resistance to the drugs (BOX 1). The total population size at each point in space (*z*) is given by 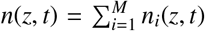, with the dynamics of the subpopulations given by

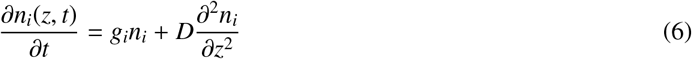

where *D* represents the common diffusion coefficient, *g*_*i*_ the growth rate of mutant *i*. For simplicity we choose

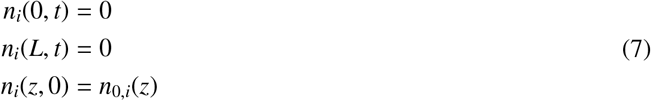

where *n*_0,*i*_ the initial distribution of mutant *i*. This system is an instance of the classical reaction-diffusion system with exponential growth kinetics, and Dirichlet boundary conditions. The assumption is that bacteria live in an idealized one-dimensional fixed domain of length *L*, and die when diffusing out the habitat, either because they meet inhospitable conditions or because of lack of resources for growth. Within the domain, we assume there are unlimited resources for growth, and there is no direct interaction between the mutants i.e. all of them are assumed to grow exponentially independently of each other.

### 2.2. Linking the model to multi-drug resistance: g_i_ = G(α_i_ x, β_i_y)

While the system 6 can be studied on its own, just as an abstract framework for mutants which vary in their growth rates and diffuse over space, here we focus on the key scenario that growth rate is entirely determined by the (α_*i*_, β_*i*_) antibiotic resistance trait of each mutant and the two drug concentrations (*x, y*) (see BOX 1). Specifically we consider two cases:

- Case 1: *g*_*i*_ = *G*(α_*i*_ *x*, β_*i*_*y*) is constant in space, i.e. a spatially *homogeneous* 2-drug environment. This leads to a constant selection coefficient everywhere in space. The frequency equation for each mutant in this case is given by: 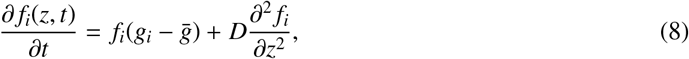 an instance of the Fisher-KPP equation (Fisher, 1937; Kolmogorov et al., 1937).
- Case 2: *g*_*i*_(*z*) = *G*(α_*i*_ *x*(*z*), β_*i*_*y*(*z*)) varies in space, i.e. a spatially *inhomogeneous* 2-drug environment, with drug concentrations that vary in *z*. This leads to a selection coefficient among mutants that is space-dependent, hence the frequency of each mutant at each point in space changes according to: 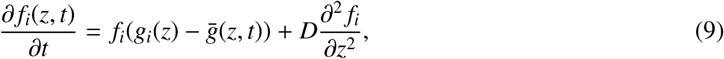

where 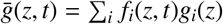

Initially we assume equal movement traits between all mutants, translating into equal diffusivities *D*, hence only growth differences driving a fitness gradient. The function *G* can be constructed analytically using abstract functional forms for antagonistic, synergistic or independent drug action (Loewe, 1953), or can be obtained empirically from growth measurement of wild-type bacteria (or an arbitrary reference strain) at a range of two-drug combination doses (*x, y*) ∈ [*x*_*min*_, *x*_*max*_] × [*y*_*min*_, *y*_*max*_]. (see, for example, (Dean et al., 2020)).

In what follows, we investigate 3 broad regimes for *G*, corresponding to i) independent drug action, ii) synergistic drug interaction, iii) antagonistic drug interaction. From Equation 9 it becomes evident that a mutant will grow in frequency at some point in space only if its growth rate is higher than the population’s mean growth at that point in space, and will decrease in frequency otherwise. Its frequency will not change if 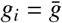.

*Multi-drug resistance evolution in space*. Combining equations above, we arrive at the following PDE system governing evolution of mean rescaling factors to drug 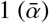, and drug 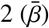), a measure of multi-drug resistance:

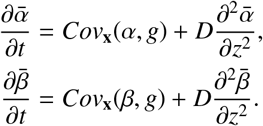

A similar link with the Price equation (Price, 1970, 1972) was derived in our earlier study (Gjini and Wood, 2021). However compared to the non-spatial model (BOX 1), in this case we are dealing with two partial differential equations, because the mean traits 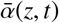 and 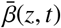 now evolve over space and time.

#### Numerical prediction of selection dynamics

While the Price Equation framework is compact, it is not by itself sufficient to predict evolution over multiple time steps since the covariance terms are dynamic. We need the explicit mutant frequencies information at a given time-point to be able to simulate the system. Thus, provided with initial conditions, i.e. an initial spatial distribution over space for all mutants, and their multi-drug resistance traits (α_*i*_, β_*i*_), a given a drug-action landscape *G* and an external drug concentration (*x*(*z*), *y*(*z*)) we can numerically integrate the equations to obtain solutions *f*_*i*_(*z, t*) for frequencies of all mutants, and finally 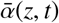 and 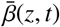as well as mean adaptation rate of the population 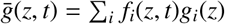.

## 3. Results

In this study, we consider the context of a cellular population adapting to a multi-drug environment via selection of pre-existing diversity. Although another route to adaptation is provided by *de-novo* mutations, we limit ourselves here to the case of standing variation, where antibiotic-resistant mutants with different degrees of susceptibility to two drugs are already present from the start, albeit at possibly very low frequencies. We investigate if and how a spatial pattern of diversity in antibiotic resistance phenotypes distribution over long time scales emerges from the fitness gradient between such mutants. We observe that a variety of selection outcomes are possible in such expanding population. We distinguish two qualitative regimes for spatial heterogeneity: i) constant drug concentration over space, and ii) spatially-varying drug concentrations, leading to constant and spatially-varying mutant growth rates over space respectively.

The main novelty of our approach is that beyond numerical simulations of the model PDEs, we propose a fully analytical measure of mutant fitness over space via which the outcome of selection can be predicted, studied and controlled.

### 3.1. An average fitness measure to predict selection outcome

When the mutants have the same growth rate everywhere in space, it is intuitive to compare them via their speed of propagation 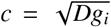, hence by their growth rates. Yet, when the growth rate is a space-dependent function, it is not clear how to establish fitness hierarchies. Our system is governed by the PDE

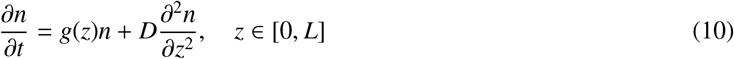

which describes each sub-population. Assuming separation of variables, and rescaling space to [0, 1] we can arrive at the following eigenvalue equation

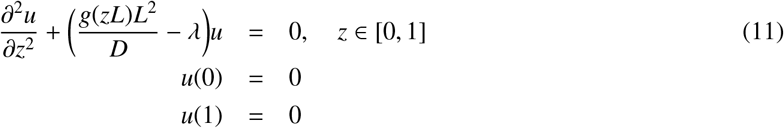

where *u* is the eigenfunction corresponding to eigenvalue λ. This falls within classical Sturm-Liouville problems, whose spectrum of equations consists of a discrete set of eigenvalues λ_*n*_ (in our case, decreasing) that determine the behavior of solutions. Importantly, when the principal eigenvalue λ_1_ > 0, it follows that the solutions grow away from zero, corresponding to the trivial spatially homogeneous steady state being unstable. These eigenvalues can be numerically computed and evaluated but they can also be analytically bounded using variational methods, or analytically approximated. The latter is the approach we adopt here.

By assuming that *g*(*z*) can be written as *g*_*max*_*g*_0_(*z*) such that ϵ≡*g*_*max*_*L*^2^/*D*≪1 is a small parameter that describes the ratio of the two relevant timescales: the timescale for diffusion (*L*^2^/*D*) and that for growth (1/*g*_*max*_, with *g*_*max*_ a suitable scaling factor, e.g. the maximum value of *g*(*z*)), the eigenvalue equation above becomes:

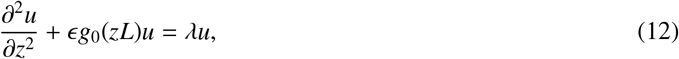

For small ϵ ≪1, it is straightforward to derive expressions for λ via classical perturbation theory (see Supplementary material S1-S3), leading to an expression for the *n*th eigenvalue λ_*n*_, up to any order in ϵ. In particular for first-order approximation we obtain:

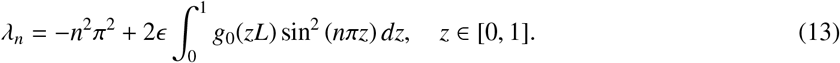

In the above expression, we can substitute ϵ explicitly. Then, reverting to original space and for *n* = 1, we obtain the following principal eigenvalue approximation:

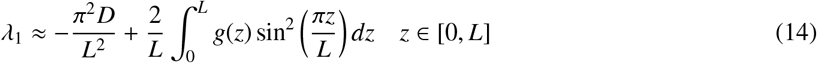

This principal eigenvalue, in the case of a single population, holds key information for successful invasion, and in the case of multiple mutants, can be used to determine their relative success in growth and propagation over space. It is noteworthy to remark that although strictly-speaking, our approximation for λ_1_ is based on assuming ϵ 1 the practical use of this asymptotic approximation typically gives reasonable results outside of its strict range of applicability. So in many numerical examples we find that the λ_1_ approximation predicts very well the outcome of selection also for small diffusion rates. However, formally the other extreme of the perturbation approach (growth much faster than diffusion) is analyzed in Supplementary material S2.

#### Special case: constant g(z) ≡ r

In the case where growth rate is a constant *r*, Equation 14 for *n* = 1 yields the survival condition λ_1_ > 0 which ultimately says that the growth rate 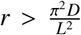, a well known result from spatial spread models (Skellam, 1951), also framed as a critical length required for the trivial steady state to become unstable 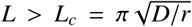. Following this reasoning, the magnitude of λ_1_, can become a relative measure by which to compare also different mutants *i* = 1, ..*M* growing and spreading in parallel. The mutant with the higher λ_1_ should win. This condition can also be applied when mutants vary in their diffusion coefficients.: the mutant with the smallest 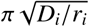 should exclude the others.

#### Arbitrary g(z)

In general, and when the population is comprised of more than one sub-population each experiencing a different space-dependent growth rate *g*_*i*_(*z*), the largest eigenvalue (Eq. 14) can be taken as a measure of average fitness over space for each mutant and we can expect that if we compare two mutants *i* and *j*, mutant *i* will ultimately dominate the population if λ_1_(*i*) > λ_1_(*j*) and vice versa. For direct comparison with simulations, we also present a discrete-space analytical approximation for λ_1_ together with some key properties (BOX 2).

##### BOX 2.

How should we define global mutant fitness over space?

To go from spatial growth rate *g*(*z*) variation among mutants to final selection outcome over space *z* ∈ [0, *L*], we propose the use of the principal eigenvalue of the linearized equation for each mutant, λ_1_. This should yield an exact measure of global fitness, which can be numerically computed or analytically approximated (see Supplementary Section S1) to obtain the mutant ranking.

###### λ_1_ when space is discretized

We begin with some suitable discretization of space, for example by subdividing the interval [0, *L*] with *K* + 2 equally spaced points (linearly-arranged patches with 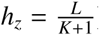). In the case where ϵ ≡ *g*_*max*_ *L*^2^/*D* is relatively small), the λ equation 14 for *n* = 1 can be approximated via secondorder centered finite differences to represent the diffusion. Written in matrix form, where also the function *g*(*z*) is now discretized as piecewise constant (*g*_*k*_) within each small sub-interval *k*, we can then use a Taylor expansion to arrive at an approximation for the principal eigenvalue λ_1_. The first-order approximation for λ_1_, under mutant growth rate *g*(*z*), for suitably high discrete resolution of space (*K* large) is given by:

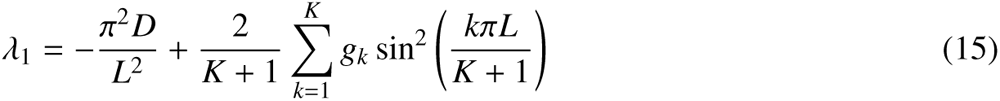

where *k* runs over all interior sub-intervals of space (See Supplementary Material S3).

###### Properties of λ_1_ as a fitness measure over space

- Under Dirichlet boundary conditions, the λ_1_ indicates that central parts of the domain have a higher value for growth than regions closer to the boundary. This can be seen from weighting of *g*(*z*) via the function sin(*z*)^2^.
- The between-mutant difference λ_1_(*i)*−λ_1_(*j*) does not depend on diffusion (first-order approximation). Diffusion rate, *D*, when equal, cannot alter selection, but it can do so when it varies, i.e. *D*_*i*_ ≠*D*_*j*_.
- In order to increase accuracy for growth functions that are very close, and enable selection sensitivity to a common diffusion rate, we must compute λ_1_ by including additional higher-order terms in the Taylor approximation (see S3.2).
- The difference λ_1_(*i*) − λ_1_(*j*) will predict the same winner as the difference in spatially-averaged growth rates 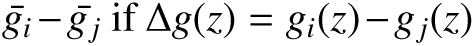 does not change sign over space *z*. This is key to compare selection under the spatial vs. non-spatial dynamics.
- Through weighting of Δ*g*(*z*) via the function sin(*z*)^2^ inside the sum (Eq. 15) and integral (Eq.14), it is evident that spatial growth function differences Δ*g*(*z*) which are symmetric around the center (*L*/2), will yield λ_1_(*j*) = λ_1_(*j*), hence coexistence between strains *i* and *j* under equal diffusion rate *D*, albeit under possible spatial segregation.

### 3.2. Example dynamics in spatially homogeneous drug concentrations

For *g*_*i*_ = *G*(α_*i*_ *x*, β_*i*_*y*), namely growth rates are independent of space, the mutant with the highest positive *g*_*i*_ wins over long time everywhere in competitive exclusion. The spatial dynamics of the frequencies of each mutant correspond to a travelling wave, with speed of spread 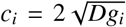. When there is no variation in *D*, the mutant with the highest positive *g*_*i*_ tends to fixation everywhere, irrespective of how this *g*_*i*_ is determined by the confluence of resistance traits and 2-drug landscape in *G*(α_*i*_ *x*, β_*i*_*y*), as illustrated in Figure 1. The only way for two (or more) mutants to coexist is their fitnesses perfectly equalize, i.e. if their antibiotic resistance traits are such that *g*_*i*_ and *g*_*j*_ fall on the same contour of *G*: *G*(α_*i*_ *x*, β_*i*_*y*) = *G*(α _*j*_ *x*, β _*j*_*y*). But the levels of such coexistence will depend on their initial total distribution, whereby the mutant with an overall advantage at the start, also persists at higher frequency in steady state coexistence.

**Figure 1:**
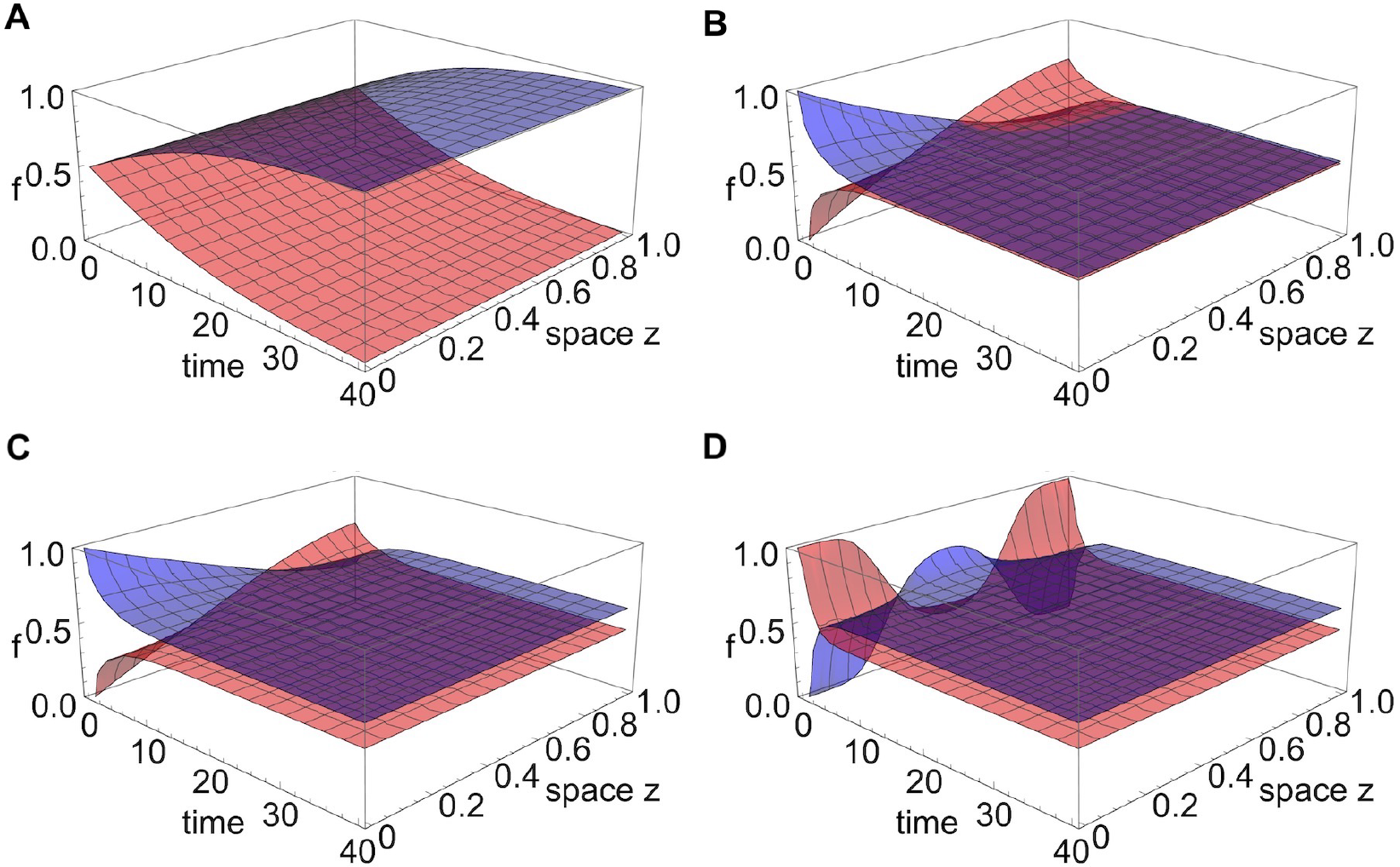
Example of outcomes among two strains for constant *g*_*i*_ over space. **A**. Competitive exclusion. In this example, strains start at uniform distribution over space, with *g*_1_ > *g*_2_, hence dynamics lead to a traveling wave solution for *f*_1_(*z, t*) and *f*_2_(*z, t*) with strain 1 traveling at faster speed and ultimately being the winner everywhere over long time. **B**. (Neutrally-stable) coexistence at 50:50 because the mutants start at equal total abundances and *g*_1_ = *g*_2_. **C**. (Neutrally-stable) coexistence different from 50:50 because mutants start at different total abundances and *g*_1_ = *g*_2_. **D**. (Neutrally-stable) coexistence different from 50:50 because mutants start at equal total abundances with *g*_1_ = *g*_2_, but their initial distribution over space favours one of them that starts at higher abundance in the center of the domain.

#### 3.2.1. Competitive exclusion when two mutants have constant g_i_ and g_j_ in [0, *L*]

Some examples of outcomes among two strains for constant *g*_*i*_ over space are illustrated in Figure 1. If these constant growth rates are different *g*_*i*_≠*g*_*j*_, under equal initial conditions, the mutant with the higher *g* will spread faster and eventually take over everywhere in space (Fig. 1A). In contrast, if these constant growth rates are equal, it is possible that asymmetric initial conditions create a bias and the mutant with a head start will eventually win (Fig. 1B-D). The bias can be created from total initial abundance (1C) or relative distribution over space (1D). In the latter case, the mutant with a relative advantage in the center of the domain will effectively spread faster and competitively exclude the other over all space.

### 3.3. Example dynamics in the presence of spatial gradients in drug concentrations

The case of *g*_*i*_(*z*) = *G*(α_*i*_ *x*(*z*), β_*i*_*y*(*z*)), when the drug concentrations can vary over space, and hence also mutant selective advantages, is the more realistic, the more interesting and naturally the more complex one. We observe mainly two results: competitive exclusion with the same mutant winning everywhere or coexistence of the same subset of mutants everywhere (although at different frequencies). The final outcome depends on how average fitness over space compares between all mutants. Analytically this PDE case is much more complex and solutions have been obtained only for special cases under certain regularities. Below we consider some specific scenarios.

#### 3.3.1. Coexistence under g(z) variation but perfect (i, j) symmetry around 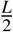

An example of linear *g*_*i*_(*z*), *g*_*j*_(*z*) variation for 2 mutants leading to coexistence everywhere is shown in Figure 2. In this case, one strain is better-adapted in the first half of the domain, the other strain is better-adapted in the second-half of the domain with the selective advantages exactly counterbalanced (Fig. 2A). For low diffusion, each strain dominates in frequency in the part of the domain where it experiences a relatively higher growth rate, maintaining a high-degree of spatial segregation in the system (Fig. 2B). As diffusion increases, the coexistence frequencies become more similar and tend towards 1/2 in both halves of the domain, leading to a more homogeneous spatial distribution of the strains over space (Fig. 2C-D).

#### 3.3.2. Competitive exclusion under broken (i, j) symmetry around 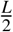, even with equal spatial averages of g

This is the example in Figure 3 A, where *g*_1_(*z*) and *g*_2_(*z*) are piecewise-constant. For equal mean growth rates over space, the situation has been long tackled analytically (Cantrell and Cosner, 1991; Seno, 1988). As recognized in previous theoretical studies, for Dirichlet boundary conditions, it is expected that the population with a spatial growth advantage in the center of the domain will experience the maximal fitness, and be the winner (Cantrell and Cosner, 1991). Indeed this is what we obtain when considering such piece-wise growth functions which make one mutant more suited to the center of the domain and the other mutant more suited to the borders of the domain. Even though mean growth rates are equal, the mutant with the central advantage spreads with an advantage and ultimately excludes the other everywhere in space.

**Figure 2:**
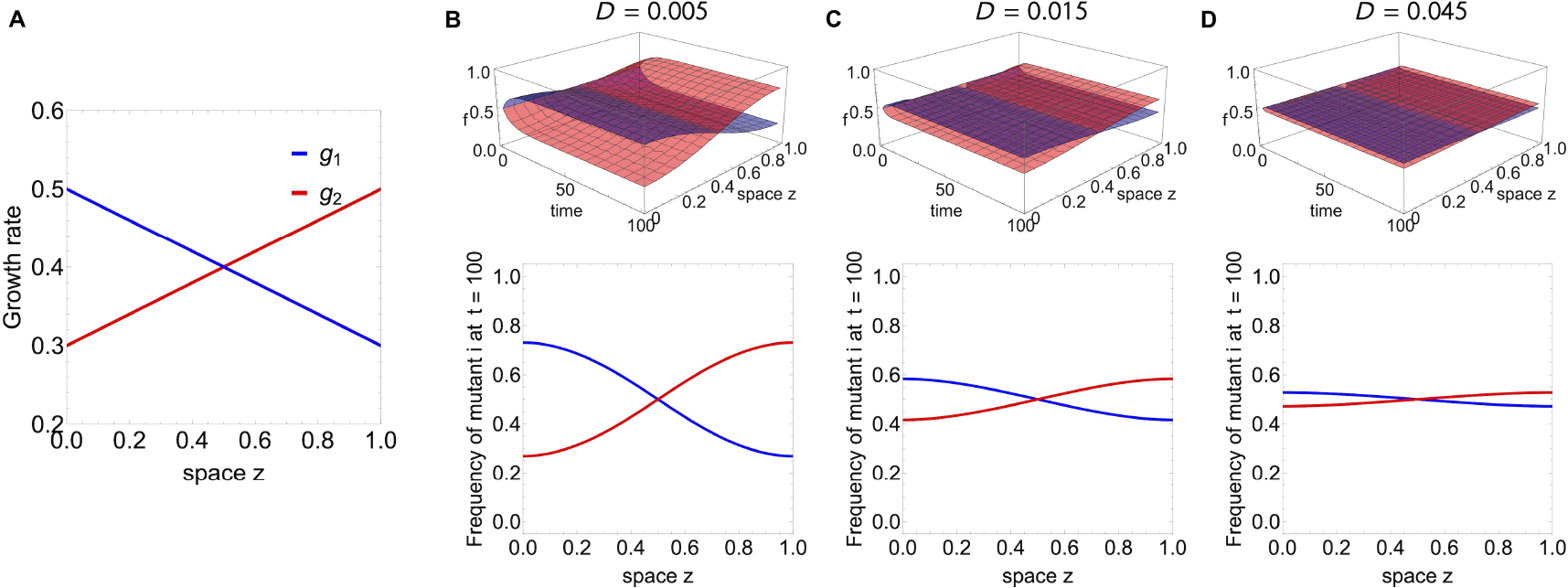
Coexistence example of 2 strains everywhere in space for space-dependent *g*_*i*_(*z*) which are mutually symmetric about *L*/2. **A**. In this case, one strain is better-adapted in the first half of the domain, the other strain is better-adapted in the second-half of the domain with the selective advantages exactly counterbalanced. **A**. For low diffusion, the two strains coexist such that each strain dominates in frequency in the part of the domain where it experiences a relatively higher growth rate, maintaining a high-degree of spatial segregation in the system. **C**. As diffusion increases, the coexistence frequencies become more similar and tend towards 1/2 in both halves of the domain. **D**. Eventually, for very high-diffusion, the growth variation starts to matter less and less, and the two strains tend to the same frequency everywhere, leading to a uniformly homogeneous spatial distribution of diversity over space.

**Figure 3:**
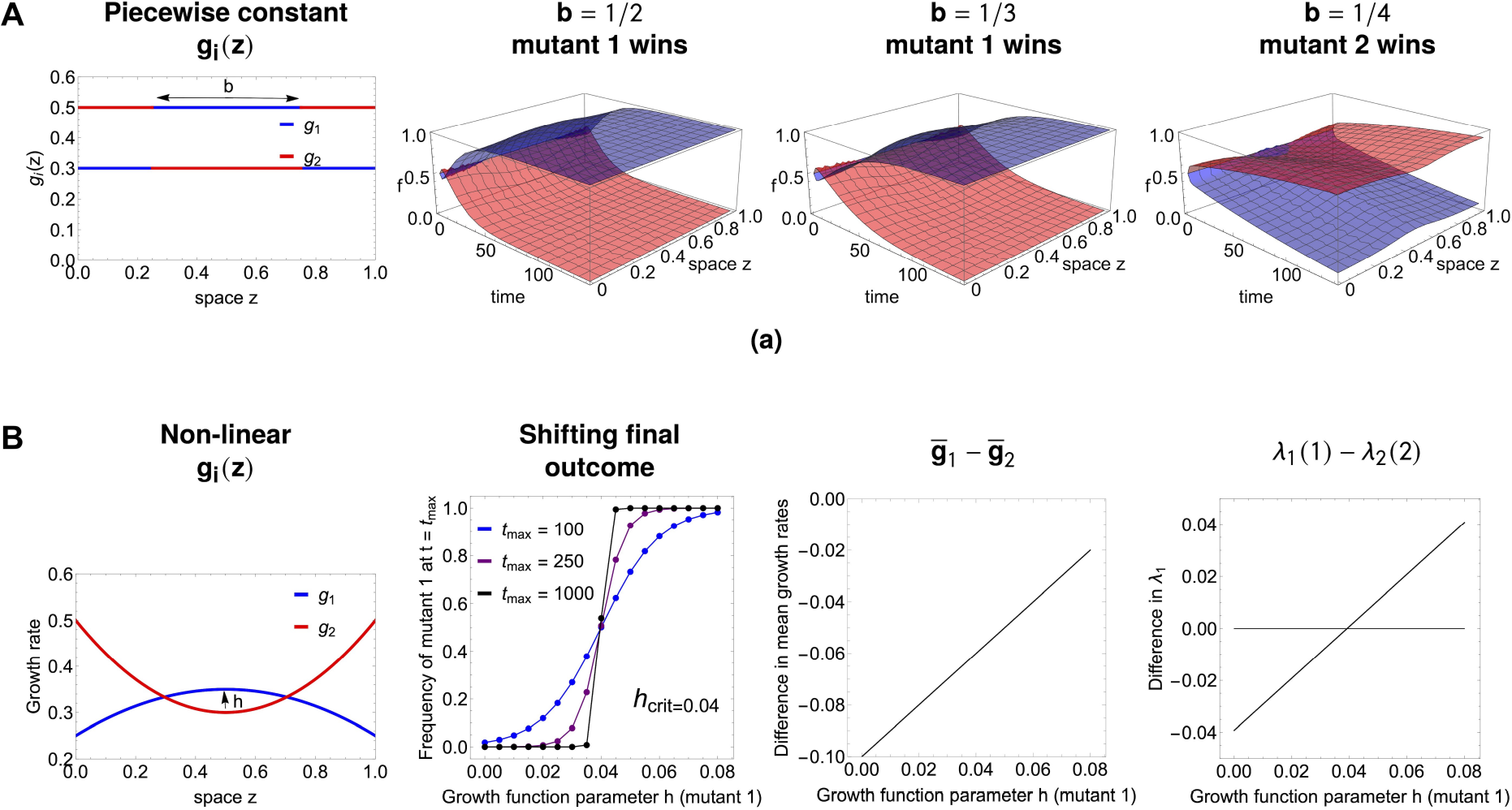
Competitive exclusion everywhere in space, but the ultimately winning strain depends on parameters of *g*_*i*_(*z*) variation. **A**. In this piece-wise growth rate example, the *g*_1_(*z*) and *g*_2_(*z*) are such that the mean growth rates for both strains are the same 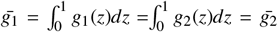 for *b* = 1/2. Yet, even with equal spatially-averaged growth rates, the strain with the central advantage will be the winner. When *b* changes, the final winner is a result of *b* as well as (*max*(*g*) −*min*(*g*)) magnitude. **B**. In this example, the winner can be overturned by modulating the width of the interval where *g*_1_(*z*) > *g*_2_(*z*), while keeping the shape of the two functions. We assume the growth rates are non-monotonic functions of space, represented by a concave and a convex parabola with vertices near the middle of the domain:*g*_1_(*z*) = *m*− σ(*z*−*L*/2)^2^ + *h* and *g*_2_(*z*) = *m* + 2σ(*z* −*L*/2)^2^ with *m* = 0.3, σ = 0.4, *D* = 0.015 and *h* varied. The critical value of *h* for overturning the final outcome is *h* = 0.04. Mutant 1 loses if *h* < 0.04 but it wins if *h* > 0.04, when its fitness advantage in the center of the domain is sufficiently high to compensate for its disadvantage near the boundaries. This cannot be predicted with the mean growth rate difference 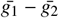 but can be predicted with λ_1_ difference for mutants 1 and 2.

In our case (Figure 3A), the spatial averages of the two mutant growth rates are simply 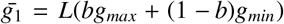 and 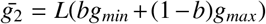, where *g*_*max*_, *g*_*min*_ are the maximum and minimum growth rate of each strain. The condition for them to be equal for any *g*_*max*_ and *g*_*min*_ is simply *b* = 1/2. In concordance with previous theory, when 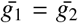, we observe that the strain with the central advantage will be the ultimate winner in the system. In contrast, the situation gets more complicated when spatially-averaged growth rates differ, hence 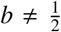 such that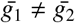. It is not always the superiority in mean *g* or in central advantage that simply drives selection. We can find cases that the strain with central advantage in *g* (in this case strain 1) may lose overall because of its total growth rate, or when the strain with superior mean *g* may lose overall because of its central fitness disadvantage. The key lies in the relative λ_1_ magnitude of each mutant.

#### 3.3.3. Tuning spatial heterogeneity of g_i_ and g_j_ can invert selection

An example with nonlinear growth rates, leading to competitive exclusion with the possibility to revert the winner via a continuous parameter change is shown in Figure 3 B. From the parabolic shapes of the growth rates it’s not immediately clear which should be the strain to ultimately grow and spread faster over space. Using λ_1_ calculations (Eq. 14) for the difference between mutants, we find that for small values of the parameter *h* it is strain 2 that ultimately excludes strain 1, but for large enough *h*, more specifically

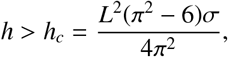

strain 1 overall fitness is superior and it will be this strain the only persisting one in the system. This result is impossible to disentangle from comparing purely spatial averages of the two growth rates. The information of the spatial average is insufficient because it weighs equally all points in space, whereas in reality the death at the boundary and the competition between diffusion and growth within the boundary makes locations near the centre of the domain be of higher value for any strain. The appropriate weighting of space is contained in the measure λ_1_.

### 3.4. An ‘atlas’ of selection outcomes under drug spatial heterogeneity: single or double resistance?

With this setup, it is now possible to systematically study scenarios of drug variation over space. First we focus just on 4 relevant mutants, which represent the main resistance combinations: mono-resistance to drug 1, and to drug 2, double intermediate resistance to both drugs, and wild-type (see Figure 4). These correspond to special locations in α, β space, namely (0, 1), (1, 0), (0.5, 0.5) and (1, 1). Then we consider synergistic vs. antagonistic drug interactions, under the assumption of a of low diffusion rate, and several explicit 2-drug concentration variation patterns over space *x*(*z*), *y*(*z*). Although more complex fitness landscape formulations are possible (Wood et al., 2014), the growth rates we assume for the drug interaction as a function of drug concentrations *x* and *y*, can be constructed for illustration, via the following simple functions:

**Figure 4:**
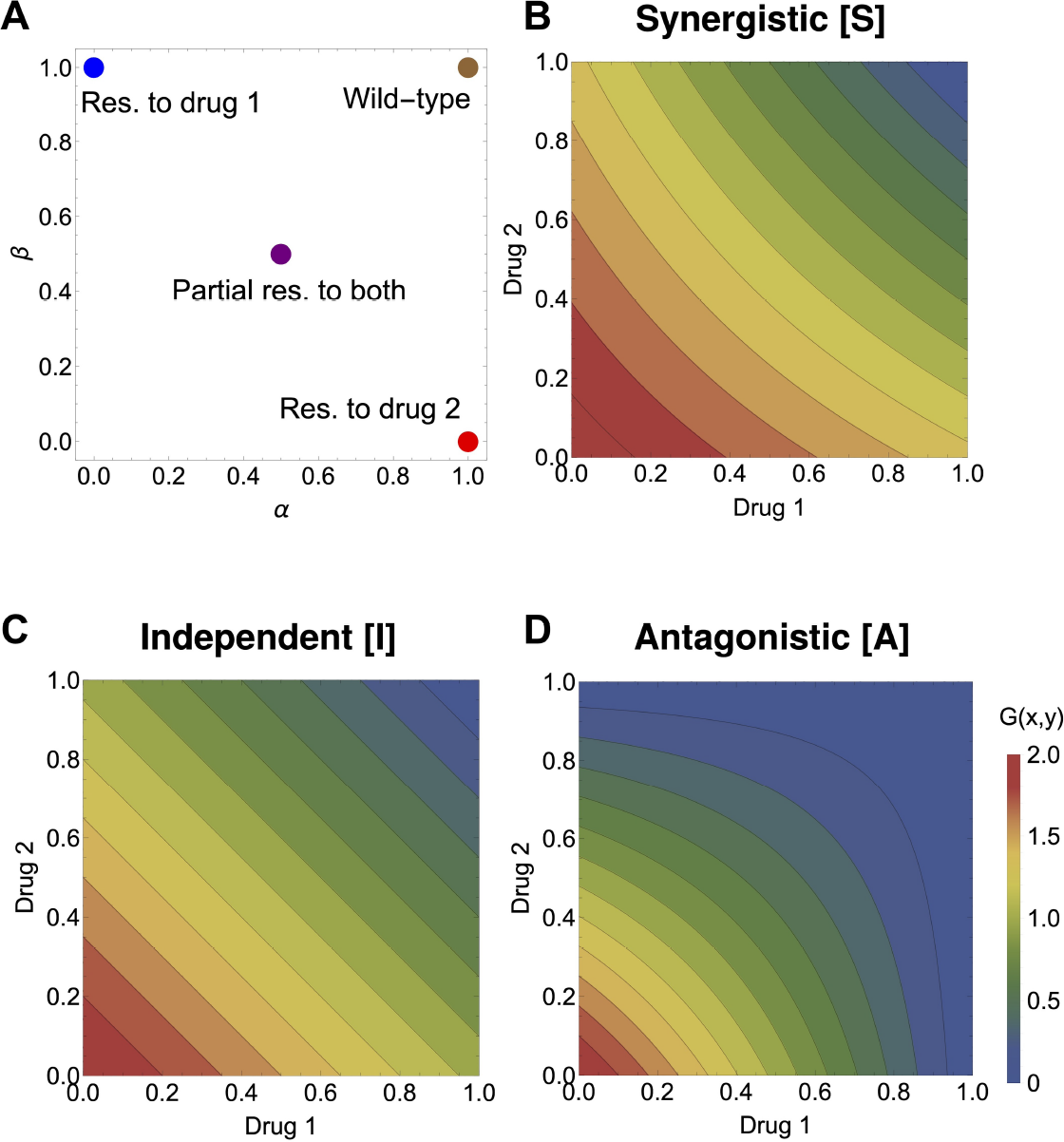
The basis for the atlas of multi-drug resistance evolution patterns over space. **A**. The four canonical mutant types for resistance phenotypes to two drugs, distributed in the (α, β) space of rescaling parameters: blue - fully resistant to drug 1 and sensitive to drug 2; red - fully resistant to drug 2 and sensitive to drug 1; purple - intermediate resistance to both drugs; brown - wild-type, sensitive to both drugs. **B**. The 3 drug fitness landscapes used: synergistic (left) independent (center) and antagonistic (right), as specified in Eqs. 16. These drug landscapes will be used to give rise to *g*_*i*_(*z*) = *G*(α_*i*_ *x*(*z*), β_*i*_*y*(*z*)) as a function of two drug variation over space *x*(*z*) and *y*(*z*). The relative fitnesses of the strains are hence dependent both on drug variation over space and on the details of the underlying growth landscape *G*.

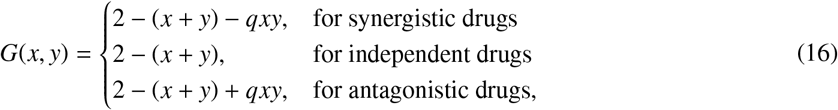

where *q* > 0 and can be taken as a measure of the strength of interaction, illustrated in Figure 4B-D.

The results of simulations are presented in Figure 5. Different multidrug concentration gradients give rise to a total of 12 scenarios, a kind of ‘atlas’. We show the frequencies obtained by simulating the system long enough for it to have reached a spatial equilibrium. These are not meant to be an exhaustive analysis but a summary of key cases of spatial heterogeneity that can shape evolution of multi-drug resistance along the main axes of monomorphic vs. polymorphic phenotypic distributions. A study of more multidrug scenarios, under a more complex growth function *G*, is shown in Figures S1 and S2-S3, where we also explore different diffusion rates.

**Figure 5:**
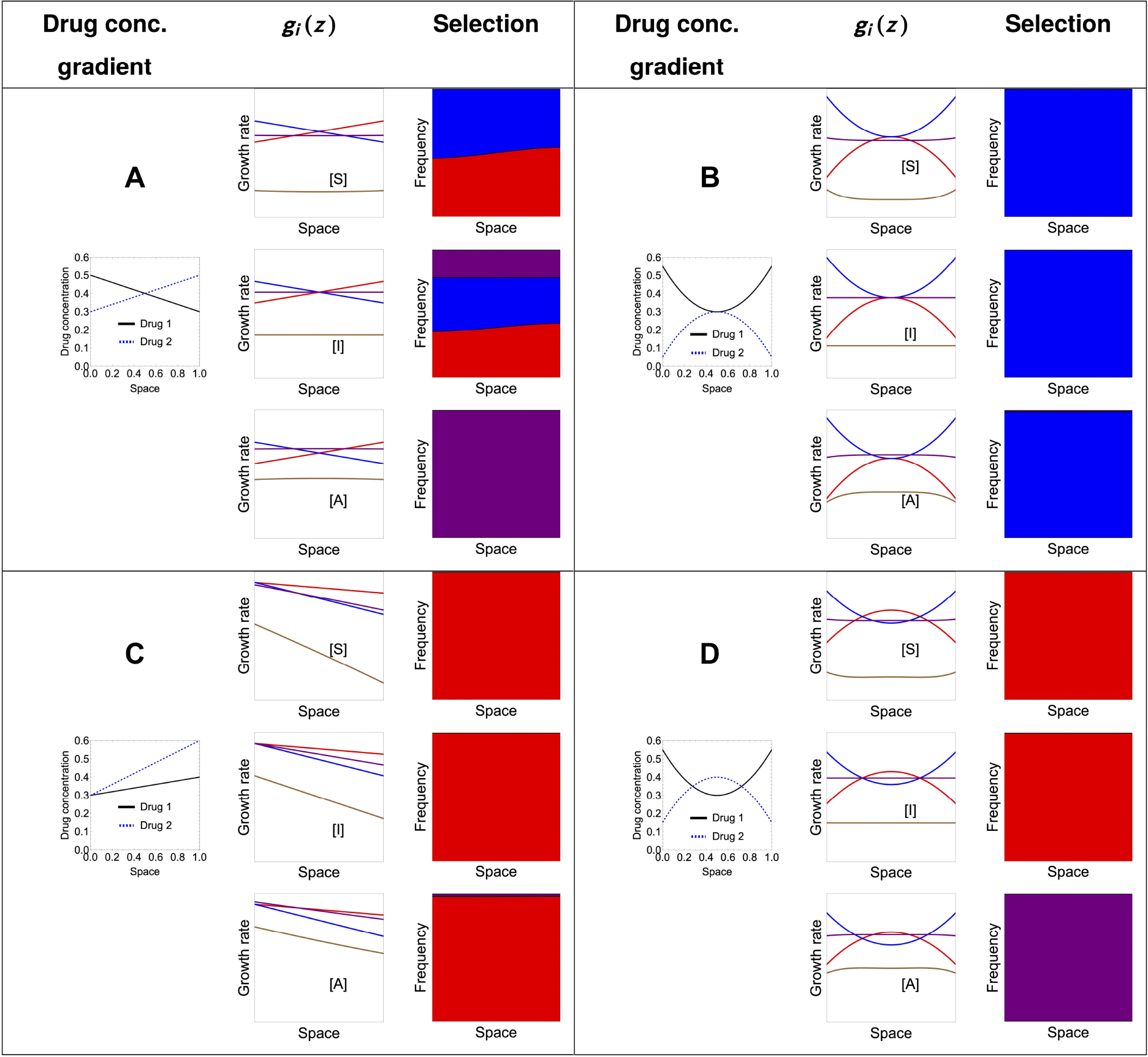
An atlas for 2-drug resistance evolution in space under spatial heterogeneity. We consider only four available mutants each with different resistance phenotypes to two drugs, distributed in the (α, β) space of rescaling parameters: blue - fully resistant to drug 1 and sensitive to drug 2; red - fully resistant to drug 2 and sensitive to drug 1; purple - intermediate resistance to both drugs; brown - wild-type, sensitive to both drugs. We considered a diffusion coefficient of *D* = 0.01; the spatial equilibrium is obtained numerically by considering the system at *t* = 1000. For all the simulations, we considered the same initial distributions with 99% wild type mutants and the remaining 1% distributed equally among the three resistant mutants. Without loss of generality, the initial distributions of each mutant were shaped as the function 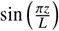, so that the homogeneous Dirichlet boundary conditions were respected. The growth landscapes were as specified in Equation 16. The interaction strength is fixed at *q* = 0.5 both in the case of synergistic and antagonistic interaction. For more drug gradient scenarios, under a more complex drug interaction profile and two diffusion rates see Supplementary Figures S1-S3.

These scenarios show that exactly anti-symmetric drug gradients relative to the center of the spatial domain are those more likely to lead to coexistence of different types of resistances (Fig.5 A), in the case of independent drugs, both mono-resistant and the double-resistant mutant coexist, in the case of antagonistic drugs, the intermediate double-resistant mutant has a higher chance to exclude the mono-resistant variants, and in the synergistic drugs regime, only the two mono-resistant mutants coexist. When one drug concentration exceed the other drug concentration throughout the domain, as intuitively expected, the mutant that gets selected in a competitive exclusion scenario uniformly over space, is the mutant that is resistant to that drug (Fig. 5B-C). When the drugs co-vary in non-linear manner over space, depending on the way this gradient translates to relative growth functions among mutants, it is possible to have different selection scenarios and fine-tune parameters to invert the hierarchical competitive fitnesses of different types of resistance mutants 5D). These selection outcomes can be entirely predicted analytically by computing and comparing the principal eigenvalues (Equation 14) between mutants in each case.

### 3.5. Verifying predictions of selection over space based on λ_1_

Next we show how the ranking based on λ_1_ comparison between mutants gives accurate prediction for final outcome of competition over space in a more complex case with more mutants (*M* > 2), more complex drug-interaction function, and arbitrary distribution of resistance phenotypes (see Figure 6).

**Figure 6:**
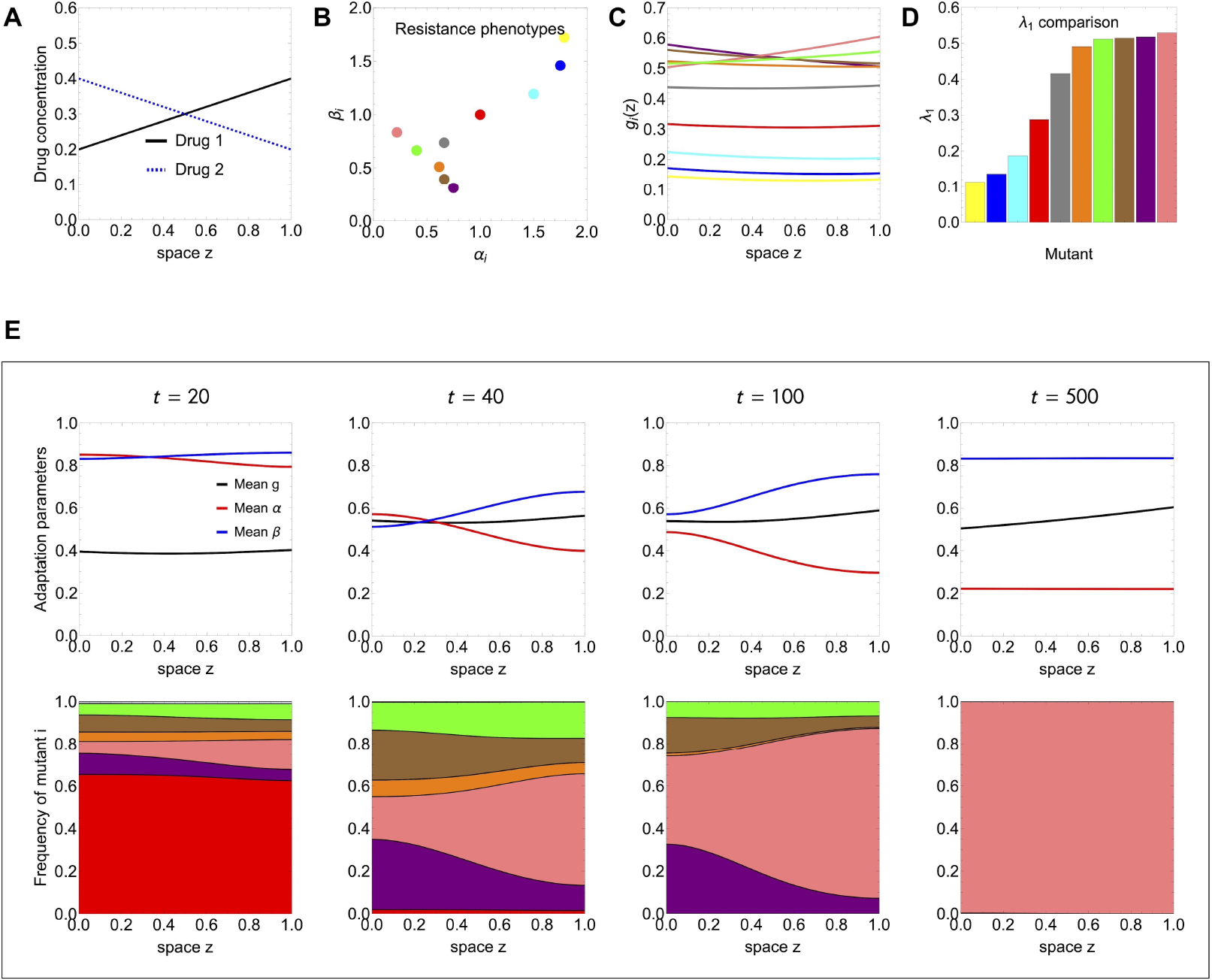
Validating selection predictions based on λ_1_ ranking among several competing mutants. We illustrate a model simulation under the linear drug gradients in A, with 10 multi-drug resistant mutants varying in (α_*i*_, β_*i*_) traits (B), growing (*g*_*i*_(*z*) in C) and spreading over space with diffusion coefficient *D* = 0.01. The λ_1_ values (Eq. 14) in D, match very well with the spatial selection dynamics observed numerically (E). Initial conditions (99% vs 1%: WT vs. all mutants) were assumed equal for all strains, satisfying the boundary conditions *n*_*i*_(*z*, 0) ∼ *sin*(π*z*).

**Figure 7:**
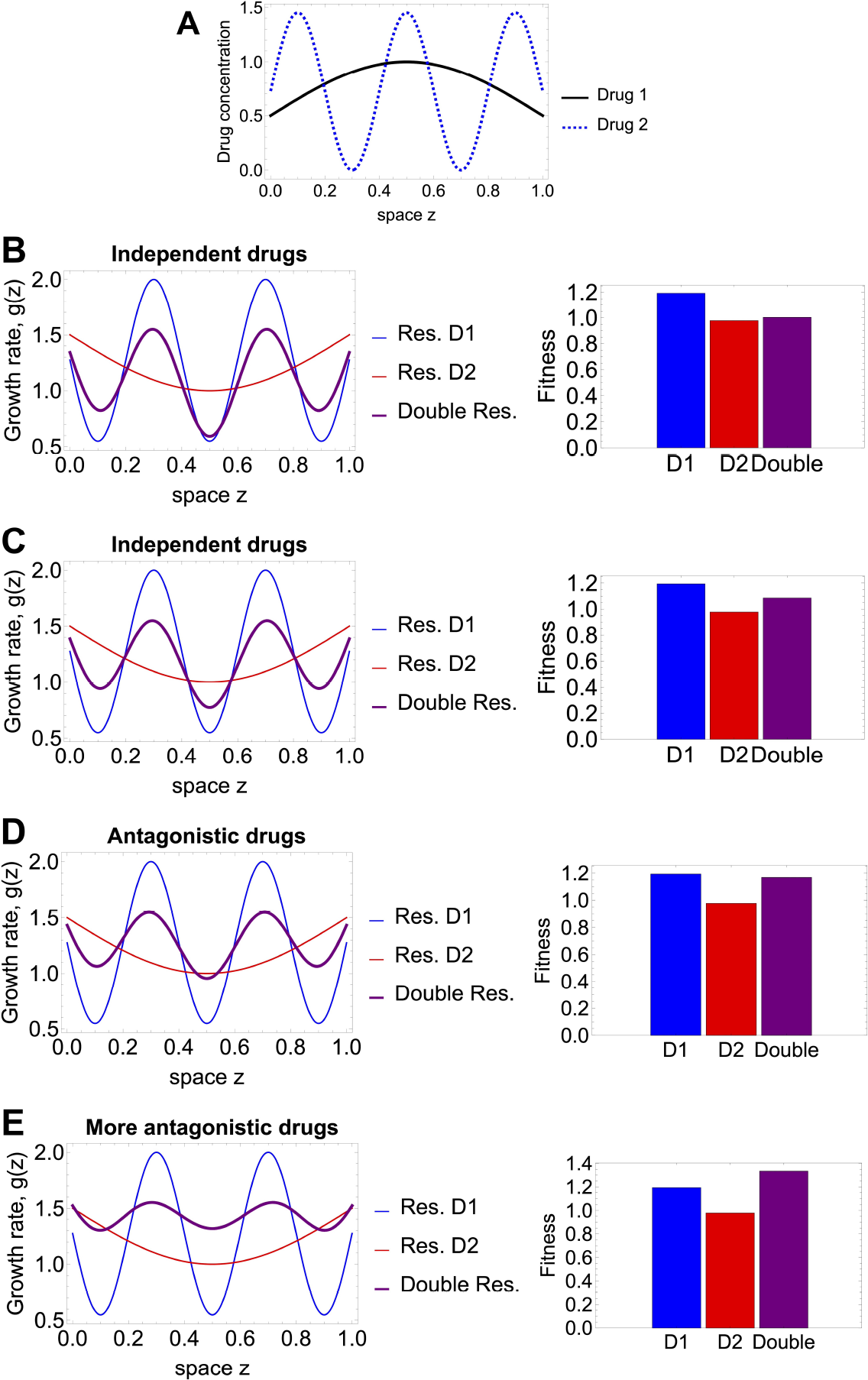
Selection outcomes for periodic drug regimes leading to periodic growth rates over space. **A**. The periodic variation of drug 1 and drug 2 over space, keeping the total amount of each drug equal. The periodic function parameters, under conservation of total drug, are: *k*_1_ = *A*_1_ = 0.5, *T*_1_ = 2 and *k*_2_ = *A*_2_ = 0.72, *T*_2_ = 0.4. Further we show mutant growth rates and selection outcome under: **B**. synergistic drug interactions; **C**. independent drugs; **C**. antagonistic drug interactions; **E**. even more antagonistic drugs. The first column shows resulting growth rates *g*_*i*_(*z*) for each mutant following the linear *G* function combinations in Eqs. 16, with *q* = 0.5 (first three rows), and *q* = 1.5 in the last row, depicting the case of stronger drug antagonism. The second column shows associated final fitnesses of the 3 mutants over space, computed on the basis of the principal eigenvalue. Assumed diffusion coefficient is *D* = 0.01.

### 3.6. Selection over space under motility and growth differences among mutants

A direct extension of this model is to, instead of assuming an equal *D*, allow for mutant-specific diffusion rates *D*_*i*_, in the environment. In this case, there is an additional trait affecting global fitness over space, namely motility. In the case of constant growth rates *g*_*i*_ it is straightforward to obtain the fittest mutant by their ranking on the classical critical criterion 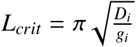 (also related to λ_1_). The mutant with the smallest *L*_*crit*_ < *L* should win. Obviously when two mutants have exactly the same fitness, perhaps by counter-balancing growth and diffusion, leading to same *L*_*crit*_ (equivalently same λ_1_) they will coexist. Whereas in the case of spatially-varying growth rates *g*_*i*_(*z*), one can resort to the same principal eigenvalue approach and compute the mutant fitness by including the assumption of a different *D*_*i*_ for each mutant. In both these cases, the diffusion traits play a key role in determining the winner or coexistence in the system, with fast diffusion sometimes being able to rescue locally maladapted strains, or slow diffusion sometimes amplifying the fitness of lower-growth variants (see Supplementary Material S4). These theoretical predictions, made accessible here through the λ_1_ comparison between variants, could be linked with existing empirical observations on bacterial coexistence and inverted competitive hierarchies driven by motility and spatial competition (Gude et al., 2020).

### 3.7. The case of periodic habitat quality: periodic multi-drug regimes

A special case of environmental variation is periodic habitat quality; in the case of multi-drug regimes, this translates to periodically-varying drug concentrations in space. This case has been long studied in the theoretical ecology literature, for example for finite one-dimensional or two-dimensional space, or infinite one-dimensional environment (Berestycki et al., 2005a,b). Typically a discretization approach is used, dividing the landscape into periodically-alternating patches of two or more types. In our case, such alternation of the space into patches of differential suitability for growth comes as a result of fitness being a direct function of the two drug concentrations. Namely for drug concentrations varying periodically in space:

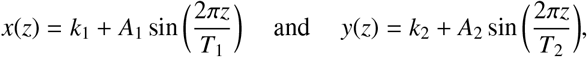

where *A*_1_ and *A*_2_ denote the amplitude of spatial variation and *T*_1_ and *T*_2_ the period of the variation for each drug, the growth rate at each point in space of the wild-type (reference strain) would be given by

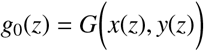

and in general, the growth rate of any variant with resistance traits (α_*i*_, β_*i*_), would be given by

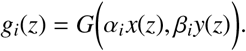

We consider again the simple drug landscapes *G* specified in Eqs.16, where with a single parameter *q* > 0, we can vary the strength of the drug interaction. Focusing on three classical mutants with traits (0, 1), (1, 0), and (0.5, 0.5), and computing their fitness based on the principal eigenvalue approach we can study which type of resistance, whether resistance to drug 1, or to drug 2 or intermediate resistance to both drugs will be favoured in each periodic drug regime, and how the result depends on the type and strength of drug interaction.

Indeed we see that for a given periodic variation of drugs over space, leading to a periodic variation of *g* over space, typically one mutant has the highest fitness. The theory predicts that under Dirichlet boundary conditions, and fixed mean growth rate over space, the population whose growth variation exhibits the highest amplitude of variation, will be the one to win (Berestycki et al., 2005a,b), e.g. the mutant who experiences a less fragmented habitat. This result is not immediately translatable to our case, since what we are controlling for is not average growth rate of each mutant, but total amount of drug 1 and total amount of drug 2.

In our simulations with given parameters, when mean growth rates of mutants may vary, we find that the mutant with resistance to drug 1 is selected in Fig.7 top panels). However, as drug interactions increase in magnitude, the intermediate double-resistant mutant can be selected (Fig.7 last panel). These outcomes of selection cannot simply be understood from just comparing the amplitude and frequency of variation in *g*(*z*), but also crucially on the mean growth rate that may change as we vary the two drugs or their interaction. In any case, computing the relative fitnesses on the basis of the principal eigenvalue ranking, leads to robust analytical results and very general predictions for whichever environmental variation, and coupling from environment to fitness.

### 3.8. Open avenues for multi-drug optimization over space

Keeping the total amount of each drug constant and equal when integrated over space, one can then try to perform optimization of periodic drug administration over space so as to select one or the other mutant. For the purposes of illustration, since detailed optimization falls beyond the scope of this paper, we studied systematically how the winning mutant varies depending on the period of a single drug’s administration over space (Figure 8), and as a function of both drugs’ spatial periods (Figure 9), in the three cases of synergistic, antagonistic and no-drug interaction. In our parameter combinations, we observe that the synergistic and independent drug actions produce very similar selection outcomes for any combination of periods of the two drugs between 0 and 2, always favouring single resistance to one drug, albeit overturning the winning strain for some critical parameter thresholds.

**Figure 8:**
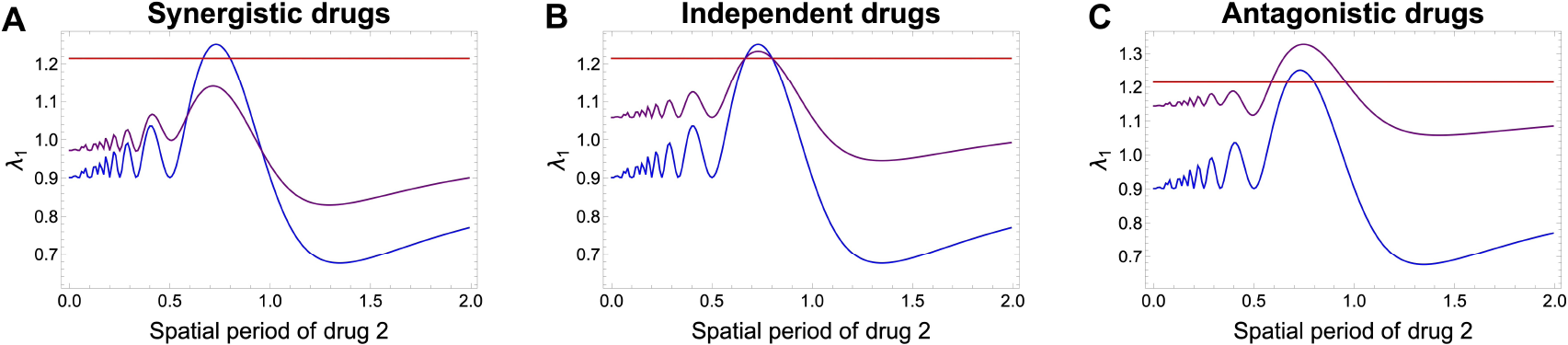
Fitness ranking among 3 mutants for periodic drug regimes, as a function of spatial period of drug 2. **A**. Synergistic drug interaction. **B**. Independent drug action **C**. Antagonistic drug interaction. The strength of interaction when assumed, was *q* = 0.5 and *G*(*x, y*) were specified as in Eq. 16. The periodic variation of drug 1 *x*(*z*) was held fixed, while drug 2 concentration *y*(*z*) over space was varied by varying the period *T*_2_. These parameters were fixed: *k*_1_ = *A*_1_ = 0.5, *T*_1_ = 0.8 and *k*_2_ = *A*_2_ = 0.5 before normalization, which then leads to total conservation of drug 1 and drug 2, fixed amount=1 for each spatial period of drug 2 *T*_2_. Final fitnesses of the 3 mutants over space, computed on the basis of the principal eigenvalue. Assumed diffusion coefficient is *D* = 0.01. In blue: single-resistance to drug 1, in red: single-resistance to drug 2, in purple: double resistant mutant with intermediate resistance to each drug.

**Figure 9:**
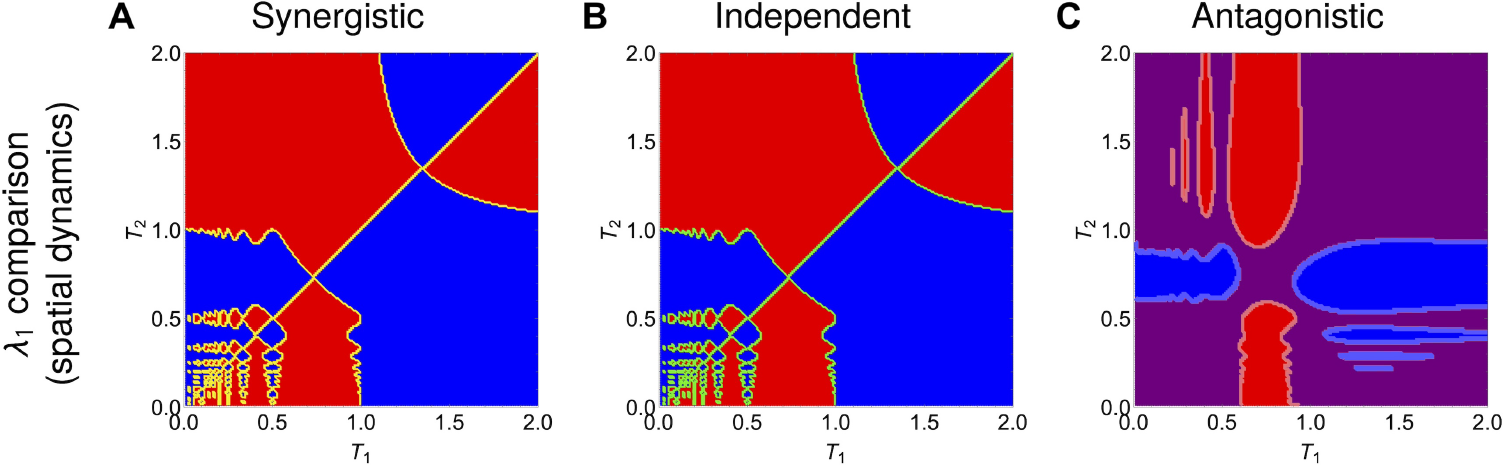
Drug-resistance selection outcomes for periodic 2 drugs as a function of their spatial periods *T*_1_ and *T*_2_. **A**. Synergistic drugs. **B**. Independent drugs. **C**. Antagonistic drugs. Shaded blue region: single-resistance to drug 1 has higher fitness, shaded red region: singleresistance to drug 2 has higher fitness, shaded purple region: double resistant mutant with intermediate resistance to each drug has the higher fitness. The periodic variations of drug 1 and drug 2 over one-dimensional space *z*∈ [0, 1] are constructed in such way as to keep the total amount of each drug equal to 1. The periodic function parameters are initially specified as: *k*_1_ = *A*_1_ = *k*_2_ = *A*_2_ for any combination of periods *T*_1_ and *T*_2_, and then immediately scaled by the integral of the periodic function over space, to obtain a total amount of drug equal to unity in each case. Assumed diffusion coefficient is *D* = 0.01. The growth functions of each mutant over space are obtained following Eqs. 16 together with the assumption that a mutant with traits (α_*i*_, β_*i*_) experiences the two drugs at concentrations α_*i*_ *x* and β_*i*_*y*. The interaction strength is fixed at *q* = 0.5 both in the case of synergistic and antagonistic interaction. In the case of synergistic/antagonistic interaction, the effect is to decrease/increase the growth rate of bacteria relative to the simple additive effect of the two drugs. See Figure S4 for the analogous figure under a more complex drug interaction function, highlighting the sensitivity to fitness landscape.

In contrast, the antagonistic drug scenario is the one that can lead also to selection of the intermediate double resistant mutant, and does so in a majority of parameter combinations. This confirms that antagonistic drug combinations, even considering spatial variation, typically constrains high-level single resistance selection, in favour of the intermediate double resistance.

Notice that two scenarios displaying the same final selection outcome only means that the ranking of the principal eigenvalues produces the same fittest strain; this does not preclude differences in the transient dynamics leading up to that outcome. Coexistence is obtained when strain fitness computed from the λ_1_ is equal. Sometimes 2 mono-resistant strains may coexist excluding the double resistance (orange border line in Fig. 9A), or the double resistant with a single monoresistant strain (region border lines in Fig. 9C), and under independent drugs, 3 strains can coexist including both mono-resistant and the double resistant strains (green border line in Fig. 9B).

In contrast, the use of the spatially-averaged growth rates in these scenarios as a proxy to predict selection would yield very different results to λ_1_. With the drug landscape defined in 16 and the total amount of the two drugs equal, we would have only one possible outcome under each drug interaction, independently of periods (*T*_1_, *T*_2_). Namely, in the synergistic case we would always have coexistence of the mono-resistance mutants, in the independent drug action case, coexistence of the three mutants (both single-resistant and the double-resistant mutant have equal 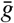), and under antagonistic drugs, the double-resistant mutant with intermediate resistance to both drugs would competitively exclude the other mutants. This result could be easily verified analytically via the integrals 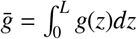 which would lead to the following growth rates:

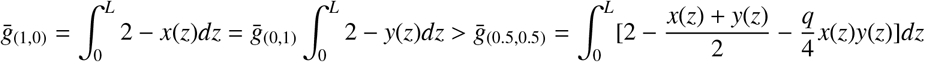

- from the assumption of total drug amount being equal 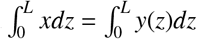, and noting *q* > 0. Hence, for whichever variation of *x* and *y* over space (i.e. independently of *T*_1_ and *T*_2_), the single-drug resistances would be selected.

The explicit analytical handle on relative fitness based on λ_1_ could further be used to design optimal multidrug regimes over space under certain constraints. For example, fixing the total amount of drug 1, and its spatial variation, which total amount of drug 2 and spatial variation would be needed to drive the double-resistant mutant toward extinction? In Figure S5 we illustrate an answer to this question, identifying precisely those drug-2 gradients over space that would be effective. Further analytical advances in multi-drug therapeutic optimization over spatially extended habitats could be obtained, by exploiting previous theoretical results Cantrell and Cosner (1991); Berestycki et al. (2005b); Pellacci and Verzini (2018) and linking them to antibiotics and bacterial realities, or generating new results on a case-by-case basis for specific microbial ecosystems under spatial gradients.

## 4. Discussion

The spatiotemporal evolution of strain frequencies in a population spreading in a homogeneous environment can be described by parallel travelling fronts where each strain propagates in space with a constant speed, according to the classical Fisher-KPP equation (Fisher, 1937; Kolmogorov et al., 1937) - an equation with a long history of study in biological invasions and population genetics (Skellam, 1951; Aronson and Weinberger, 1978). Under the classical exponential or logistic growth kinetics, the result of such competition typically leads to competitive exclusion where the fittest strain will be the only one to survive everywhere in space over long time.

In a spatially-varying environment, the local quality of the habitat affects the speed of spread of an invading population. Natural environments where populations grow and spread are generally heterogeneous, composed of different sub-habitats, such as forests, plains, marshes, and the like, or consist of pieces divided by barriers such as roads, rivers, cultivated fields (Kinezaki et al., 2003). This case has a long history of ecological, environmental and agricultural interest, and a long history of mathematical study with analytical results on periodic traveling waves (Shigesada et al., 1986), piecewise-environmental variation (Cantrell and Cosner, 1991) and up to the more recent work by (Berestycki et al., 2005a,b) on periodic spatial variation, with the pulsating front characterized by its average speed.

The key quantity highlighted in many of these earlier studies is the principal eigenvalue of the linearized equation, which determines the global stability of the stationary state 0. Although these studies were primarily interested in biological invasion of a single species, such result can be applied to the context of multiple strains of a population spreading in a heterogeneous habitat. One can use the same logic for ranking the fitness of different mutants on such heterogeneous habitat, or alternatively for determining relative environmental suitability for each mutant. Ranking the global (in)stability of the 0 steady state, via principal eigenvalue comparisons, allows to reach a conclusion about the strains’ survival in an environment that is differentially suitable, and hence predict selection outcomes in the system over long time.

Harnessing the analytical foundations of these results, we go here one step further by applying this theory to the context of antibiotic resistance evolution, and providing an accurate approximation of this principal eigenvalue (fitness measure). We study arbitrary variation in the environment suitability, linking spatial heterogeneity to explicit multi-drug antibiotic regimes, and integrating fitness landscapes, drug interactions and collateral effects, with the aim to predict multi-drug resistance evolution as a selection process among any mutants. Other studies have considered the role of spatial heterogeneity in the evolution of resistance, using the framework of an epidemiological model and focusing on the case of a single drug being used with periodic variation in the growth rates of single and double-resistant genotypes (Griette et al., 2022). Here we develop a more general and comprehensive link between traveling fronts and multi-drug resistance. In our framework, periodic variation can be seen as a special case of spatial heterogeneity, when drugs are used at periodic concentrations over space. Furthermore, differently from (Griette et al., 2022), we include the possibility of multiple drugs interacting, which affects mutant success and final competitive outcome between multi-drug resistant variants.

The main advantage of this framework is that it provides a simple and general template for studying multi-drug resistance evolution in space, possibly applicable to other systems and open to analytical extensions (BOX 3). In particular, it allows for continuous resistance traits, includes collateral effects and drug interactions explicitly, and the prediction of final outcomes is based on dominant eigenvalue ranking among mutants, which can be analytically approximated. The framework is easily extendable to include spatial variation of multiple drugs to > 2 drugs, and hence enabling study of higher-dimensional evolution in antibiotic resistance fitness traits. Especially in the case of 1-d and 2-d environments, there are many results that can be directly applied from the literature to the case of antibiotic resistance, such as optimal spatial variation to prevent or facilitate global spread of an invading species or strain (Cantrell and Cosner, 1991; Berestycki et al., 2005a).

Mathematically, and strictly-speaking the λ_1_ presented in our framework corresponds to a growth rate, related to the negative principal eigenvalue in other studies (Cantrell and Cosner, 1991; Berestycki et al., 2005a; Pellacci and Verzini, 2018), whose optimization for invasion and persistence would seek a minimum. While this is a convention, our technical choice enables us conceptually and practically to rank the mutants more easily favouring the one with the relatively higher λ_1_. Other approaches to compute or obtain suitable bounds for λ_1_ can come from variational methods applied to Sturm-Liouville problems and the Rayleigh quotient (Cantrell and Cosner, 1989). We adopted a perturbation theory approach, which despite its requirement for ϵ small, appears to apply well outside this immediate strict range. In particular, the λ_1_ approximation matches very well the selection results that the model simulations display even in cases of lower diffusion.

A disadvantage of our approach is that while the population involved and its constituent strains are considered dynamic, the environment, namely the drug concentrations across space, is generally regarded as static. Other modeling frameworks are needed to treat situations with both temporal and spatial variation in the environment, or a mutual feedback, such as in resource-based models where cells physically interact with resources at the expanding front, e.g. in biofilms (Young and Allen, 2022; Sinclair et al., 2019). Similarly, by assuming large populations, we neglect stochastic fluctuations in sub-populations which might affect selection outcomes in certain settings.

### BOX 3.

General applicability of the framework, outlook and challenges

#### Spatial growth and selection under multiple stressors

The simple framework based on rescaling parameters (α, β) could be applied to other biological populations, at different scales, growing in response to multiple stressors and spreading through migration in heterogeneous environments that generate a gradient for growth. Differential variant makeup, whether genetic or non-genetic, but heritable, that produces different susceptibility traits to these stressors in the population forms the basis for selection on relevant timescales, manifested in its simplest form as competitive exclusion (*monomorphism*) or coexistence patterns (*polymorphism*) in spatially extended habitats. The stressors could range from antibiotics (Larsson and Flach, 2022), to agrochemicals (Malagón-Rojas et al., 2020), temperature, moisture, (Jiang et al., 2017), pH, salinity (Wicaksono et al., 2022), to nutrient levels, oxygen, or physiological micro-environments (Chikina and Vignjevic, 2021). The primary dose-responses of *stressor-to-growth* phenotypic effect can be used to obtain (α_*i*_, β_*i*_) traits in the sub-populations of interest, and then rescale accordingly the *G*(*x, y*) of the reference (WT) strain to obtain growth rates *g*_*i*_ for all variants under any combination of stressor concentrations.

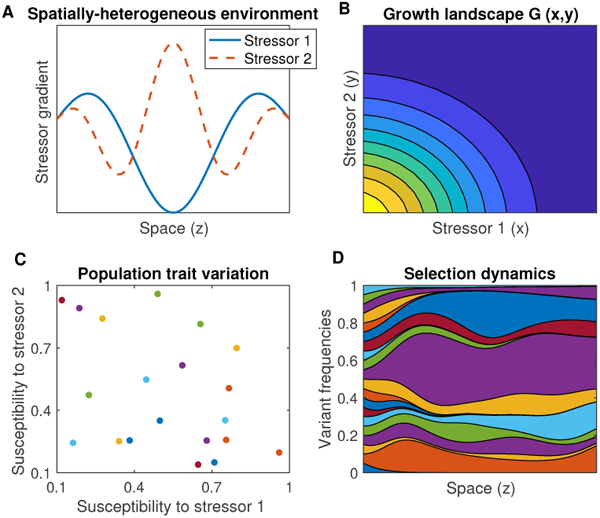

#### Limitations and caveats

In some cases, scaling factors may not fully capture the variation in mutant growth relative to wild-type as a function of stressor concentrations. Much more nonlinear functional transformations may be required to obtain mutant fitness landscapes, and this remains an active area of research (Wood et al., 2014). Similarly, the exponential model may be too unrealistic to represent the intricate mechanisms of growth and interactions between strains (Maciel and Lutscher, 2018; Estrela and Brown, 2018), thusfar assumed negligible. We focused on the case of mainly positive growth rates. However, locally negative growth rates (e.g. supra-inhibitory stressor doses) could complicate outcomes, via stronger dependence on initial conditions, or additional sensitivity to variable diffusion rates to compensate for fitness troughs.

#### Extensions and outlook

Applications can be envisaged in other systems such as gut microbiota, soil bacteria and their spatio-temporal distribution under abiotic gradients, cancer cell populations and drug resistance selection dynamics along physiological gradients, antibiotic resistance evolution at the epidemiological scale, toxicology data, freshwater aquatic systems and environmental biotechnology. More than 2 stressors could be implemented, thereby yielding high-dimensional susceptibility traits. An additional axis of resistance cost can be integrated, either as independent or via a functional dependence on α, β traits. The model could include space-dependent diffusion or habitat preference bias at the interface between distinct environmental patches (Maciel and Lutscher, 2018). Analytical extensions could exploit links with homogenization techniques from landscape ecology (Yurk and Cobbold, 2018) and global fitness perspectives from adaptive dynamics (Metz et al., 1992).

We only included diffusion, i.e. random movement of cells in space, while other similar reaction-diffusion models focusing on microbiota composition variation along the gut have included both diffusion and directed flow of bacterial lineages along a longitudinal growth gradient (Ghosh and Good, 2022). We also did not explicitly include active mutation processes in the kinetics of the spatial model, a process which would break the independence between the existing strains (Gjini and Wood, 2021), and preclude the straightforward application of the first eigenvalue approach for fitness comparison. In the particular case in which mutation rates to a given variant are equal among all possible parental strains, one could plausibly assume the initial total distribution among mutants as a proxy for such hierarchical mutation biases, and apply still the present model. *De-novo* diversity generation could be added with specific assumptions on parent and offspring phenotypes, dependence on the current environment, population size, and/or spatially-varying mutation rate. This remains an interesting avenue for the future.

Another mechanism to break the independence between sub-populations would be competition or facilitation between variants, explicitly embedded into their growth kinetics and spatial spread. Other studies have addressed it e.g. in ecological models (Maciel and Lutscher, 2018; Estrela and Brown, 2018) or epidemiological multi-strain systems (Le et al., 2023; Le and Madec, 2023), leading to potentially very complex replicator-type dynamics. Although we did not focus on analytical results for coexistence levels between strains, when their global fitnesses equalized over space, these results can be easily obtained from the same approximation steps based on perturbation, that allowed us to compute the dominant eigenvalue for each mutant (see S1-S3). On the other hand, analytical results for total population size and frequencies could be more easily obtained for cases constant growth rates over space, applying classical reaction-diffusion theory and travelling-wave characterization of the solutions.

While the debate of antibiotic resistance management (Raymond, 2019) has optimization at the center, we very briefly sketched a few aspects of the model to inform optimal combination multi-drug therapies over space (e.g. Fig. 9 and S5). Analytical results on optimization remain challenging even for simple piecewise growth variation as recognized in earlier work (Cantrell and Cosner, 1991). Many factors do play a role, including sensitivity to boundary conditions and structure of habitat fragmentation (Berestycki et al., 2005a; Pellacci and Verzini, 2018), modelling assumptions, nonlinearities linking phenotypes to growth and variation in habitat ‘quality’, and details of underlying stressor interaction. However, the general framework presented here for multi-drug gradients can be a basis that can be tailored to specific systems and their optimal control in the future. We expect many opportunities for model-data links both in microbiology and lab evolution experiments, as well as in the larger-scale epidemiology of multi-drug resistance evolution.

## Funding

TF was supported by grant GL Proj.2022/0006 awarded by Fundacão Luso-Americana para o Desenvolvimento (FLAD). EG acknowledges support by the FCT (CEECIND/03051/2018). KW is supported by NIH R35GM124875.

## Supporting information

Supplementary Notebooks and Movies

## Notes

### Competing Interest Statement

The authors have declared no competing interest.

