## Supplementary figures and images for "Modeling spatial evolution of multi-drug resistance under drug environmental gradients"

### pinkwins2.gif

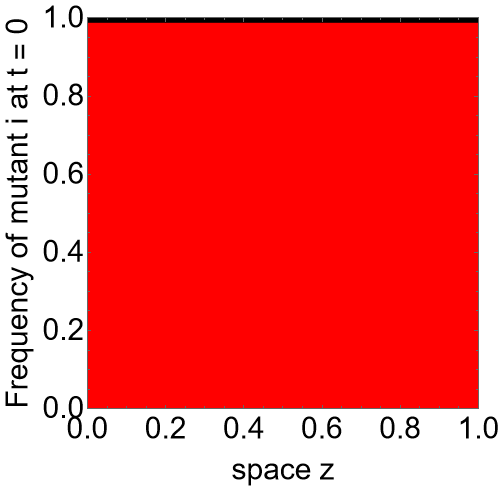

### purplewins2.gif

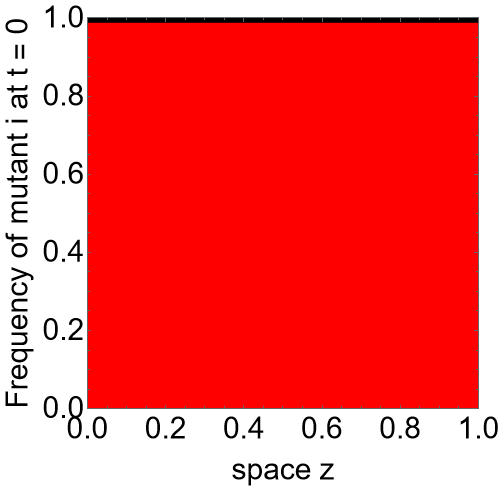
