## Supplementary Material for "Modeling spatial evolution of multi-drug resistance under drug environmental gradients"

### Supplementary Information

#### S1. Using perturbation theory for continuous space models to derive $\lambda_1$

We start from the original PDE equation

$$\frac{\partial n(z,t)}{\partial t} = D\frac{\partial^2 n(z,t)}{\partial z^2} + g(z)n(z,t) \tag{S1}$$

To understand the behavior of solutions, let's assume separation of variables, that is that $n$ can be written as a product of two functions $T$ and $S$, of time and space only:

$$n(z,t) = S(z)T(t)$$

Then the PDE can be written as:

$$\dot{T} = DS''T + g(z)ST \tag{S2}$$

where $S''$ refers to second-order spatial derivative of $S$. In this equation we can gather terms in the following way, and set the left-hand side and right-hand side of the equation to a constant $\lambda$:

$$\frac{1}{D}\frac{\dot{T}}{T} = \frac{S''}{S} + \frac{g(z)}{D} = \lambda \tag{S3}$$

So we end up with a system of equations for $T$ and $S$:

$$\frac{dT}{dt} = \lambda DT \tag{S4}$$

$$\frac{d^2S}{dz^2} + \frac{g(z)}{D}S = \lambda S \tag{S5}$$

where $z \in [0, L]$ and $S(0) = 0$ and $S(L) = 0$. The latter gives us the eigenvalue equation for $\lambda$:

$$\boxed{\frac{d^2S}{dz^2} + \left(\frac{g(z)}{D} - \lambda\right)S = 0.}$$

*Rescaling space to* $[0, 1]$

Let's rewrite the eigenvalue in a new spatial coordinate $z_{new} = z/L \in [0, 1]$:

$$\boxed{\frac{d^2S(z_{new})}{dz_{new}^2} + \left(\frac{L^2 g(z_{new}L)}{D}\right)S(z_{new}) = \lambda S(z_{new}), \quad z_{new} \in [0,1]}$$

or more compactly:

$$\boxed{\frac{d^2S}{dz_{new}^2} + \left(\frac{L^2 g(z_{new}L)}{D}\right)S = \lambda S, \quad z_{new} \in [0,1]} \tag{S6}$$

In general, the above should be a regular Sturm-Liouville problem, and we should be able to numerically compute its eigenvalues. Consider

$$\frac{L^2 g(z_{new}L)}{D} = \frac{L^2 g_{max}}{D} g_0(z_{new}L)$$

*Case 1: $\frac{L^2 g_{max}}{D}$ relatively small, $\epsilon = \frac{L^2 g_{max}}{D} \ll 1$.* In the case of high-diffusion relative to the growth term above, we may approximate the spectrum of these equations to obtain the eigenvalues. Simplifying the notation to use $z$ for the new variable $z_{new} \in [0,1]$ and $u(z)$ for $S(z_{new})$, we have the following

$$\frac{\partial^2 u(z)}{\partial z^2} + [\epsilon g_0(zL) - \lambda]u(z) = 0, \tag{S7}$$

Thus, we need to solve an equation of the form

$$\frac{\partial^2 u(z)}{\partial z^2} + \epsilon g_0(zL)u(z) = \lambda u(z), \quad z \in [0,1] \tag{S8}$$

subject to boundary conditions $u(0) = u(1) = 0$. The function $g_0(zL)$ is of order unity and describes growth over space, $\epsilon$ is a dimensionless parameter, and $\lambda$ ultimately will determines the stability of the 0 steady state.

***A simpler problem.*** We first start with a simpler problem defined by

$$\mathcal{F} y_n(z) = \sigma_n y_n(z) \tag{S9}$$

where $\mathcal{F} \equiv \frac{\partial^2}{\partial z^2}$ is the diffusion operator, $y_n(z)$ is the nth eigenfunction, and $\sigma_n$ is the nth eigvenalue of the operator. In this case, the (normalized) eigenfunctions and eigenvalues that satisfy the boundary conditions are given by

$$y_n(z) = \sqrt{2}\sin(n\pi z), \tag{S10}$$

and

$$\sigma_n = -n^2\pi^2, \tag{S11}$$

with $n = 1, 2, 3, ....$ These eigenfunctions form an orthonormal basis for functions on our domain. In what follows, we will use Dirac's bra-ket notation, so (for example) Equation S9 becomes

$$\mathcal{F}\,|y_n\rangle = \sigma_n\,|y_n\rangle \tag{S12}$$

and our problem is to solve Equation S8, which becomes an eigenvalue equation of the form

$$(\mathcal{F} + \epsilon W)\,|u_n\rangle = \lambda_n\,|u_n\rangle\,. \tag{S13}$$

where $W = g_0(zL)$. In the limit $\epsilon << 1$, we can use perturbation theory to approximate $\lambda$. This problem is analogous to one that arises in Quantum Mechanics when the Hamiltonian (analogous to $\mathcal{F}$, for which the eigenvalue problem is solved) is perturbed by a small perturbation, in this case $\epsilon W$ Sakurai and Commins (1995). Briefly, we expand the solution $|u_n\rangle$ and the eigenvalue $\lambda_n$ as power series in $\epsilon$,

$$\begin{aligned} |u_n\rangle &= |u_n^0\rangle + \epsilon\,|u_n^1\rangle + \epsilon^2\,|u_n^2\rangle \ldots \\ \lambda_n &= \lambda_n^0 + \epsilon\lambda_n^1 + \epsilon^2\lambda_n^2 \ldots \end{aligned} \tag{S14}$$

where subscripts label the nth eigenvalue or eigenfuction and subscripts indicate the term in the series expansion. Plugging these expressions into Equation S13 yields, at order $\epsilon^0$,

$$\mathcal{F}\,|u_n^0\rangle = \lambda_n^0\,|u_n\rangle\,, \tag{S15}$$

so to zeroth order the solution is that of the unperturbed problem and is given above (Equations S10 and S11). To order $\epsilon^1$, we have

$$\mathcal{F}\,|u_n^1\rangle + W\,|u_n^0\rangle = \lambda_n^0\,|u_n^1\rangle - \lambda_n^1\,|u_n^0\rangle\,. \tag{S16}$$

If we take the scalar product of the above equation with the unperturbed solution, we get

$$\langle u_n^0|\,\mathcal{F}\,|u_n^1\rangle + \langle u_n^0|\,W\,|u_n^0\rangle = \lambda_n^0\,\langle u_n^0|u_n^1\rangle + \lambda_n^1\,\langle u_n^0|u_n^0\rangle\,. \tag{S17}$$

Because $\mathcal{F}$ is self-adjoint with these boundary conditions–which one can prove using integration by parts twice–the first term on the left side is identical to the first term on the right side of Equation S17. Using the orthonormality of the unperturubed solutions, we are left with

$$\lambda_n^1 = \langle u_n^0 | W | u_n^0 \rangle = 2 \int_0^1 g_0(zL) \sin^2(n\pi z)\, dz. \tag{S18}$$

So the expression for $\lambda_n$ to order $\epsilon$ is given by

$$\lambda_n = -n^2\pi^2 + 2\epsilon \int_0^1 g_0(zL) \sin^2(n\pi z)\, dz, \quad z \in [0, 1] \tag{S19}$$

Substituting $\epsilon$ this becomes:

$$\lambda_n = -n^2\pi^2 + 2\frac{L^2 g_{max}}{D} \int_0^1 g_0(zL) \sin^2(n\pi z)\, dz, \quad z \in [0, 1] \tag{S20}$$

If we return to original units of time and space, this equation becomes

$$\lambda_n = -\frac{n^2\pi^2 D}{L^2} + \frac{2}{L} g_{max} \int_0^L g_0(z) \sin^2\left(\frac{n\pi z}{L}\right) dz, \quad z \in [0, L]. \tag{S21}$$

In particular for $n = 1$ and recalling $g(z) = g_{max} g_0(z)$, we have:

$$\boxed{\lambda_1 = -\frac{\pi^2 D}{L^2} + \frac{2}{L} \int_0^L g(z) \sin^2\left(\frac{\pi z}{L}\right) dz, \quad z \in [0, L].} \tag{S22}$$

The special case of the principal eigenvalue ($n = 1$ ) when the growth rate is constant over space, namely $g(z) \equiv r$, we have:

$$\lambda_1 = -\frac{\pi^2 D}{L^2} + \frac{2}{L} r \int_0^L \sin^2\left(\frac{\pi z}{L}\right) dz = -\frac{\pi^2 D}{L^2} + \frac{2}{L} r \frac{L}{2} = -\frac{\pi^2 D}{L^2} + r. \tag{S23}$$

And

$$\lambda_1 > 0 \iff L > \pi \sqrt{\frac{D}{r}}$$

ensures growth of the population, a classical result in reaction-diffusion models with exponential kinetics, known as the critical patch size.

We can also write down an expression for $|u_n\rangle$ to first order in $\epsilon$ as

$$|u_n\rangle = |u_n^0\rangle + \epsilon \sum_{m \neq n} \frac{\langle u_n^0 | W | u_m^0 \rangle}{\lambda_n^0 - \lambda_m^0} |u_m^0\rangle = \sqrt{2} \sin(n\pi z) + \epsilon \sum_{m \neq n} \frac{2 \int_0^1 g_0(z) \sin(m\pi z) \sin(n\pi z)\, dz}{\pi^2 (m^2 - n^2)} \sqrt{2} \sin(m\pi z) \tag{S24}$$

<u>*Higher order approximations.*</u> Using a similar strategy, one can show that at order $\epsilon^k$, the approximation becomes

$$\lambda_n^k = \langle u_n^0 | W | u_n^{k-1} \rangle. \tag{S25}$$

At order $\epsilon^2$, we have

$$\lambda_n^2 = \langle u_n^0 | W | u_n^1 \rangle = \sum_{m \neq n} \frac{\left(\langle u_m^0 | W | u_n^0 \rangle\right)^2}{\lambda_n^0 - \lambda_m^0} \tag{S26}$$

which reduces to

$$\lambda_n^2 = \sum_{m \neq n} \frac{4 \left[\int_0^1 g_0(z) \sin(m\pi z) \sin(n\pi z)\, dz\right]^2}{\pi^2 (m^2 - n^2)}. \tag{S27}$$

#### S2. WKB approximation in the slow diffusion limit $\frac{D}{L^2} \ll g_{max}$

Starting from the eigenvalue equation S6

$$\frac{\partial^2 u(z)}{\partial z^2} + \left[\frac{g(z)L^2}{D} - \lambda\right]u(z) = 0, \quad z \in [0,1] \tag{S28}$$

if we are in the opposite regime of low diffusion relative to the growth rate, we can rewrite the above as:

$$\frac{D}{L^2 g_{max}} \frac{\partial^2 u(z)}{\partial z^2} + [g_0(z) - \lambda]u(z) = 0, \tag{S29}$$

where $g_0(z) = \frac{g(z)}{g_{max}}$ is the rescaled growth function by factor $g_{max}$. In this way, our eigenvalue problem becomes

$$\epsilon \frac{\partial^2 u(z)}{\partial z^2} + [g_0(z) - \lambda]u(z) = 0, \tag{S30}$$

where the small perturbation parameter is given by:

$$\epsilon = D/(g_{max}L^2) \ll 1.$$

Thus we can use the WKB approximation–a type of multi-scale perturbation theory Sakurai and Commins (1995)–to approximate Equation S30. Specifically, we assume a solution of the form

$$u(z) \propto \exp\left[\delta^{-1}\left(S_0(z) + \delta S_1(z) + \delta^2 S_2(z)...\right)\right] \tag{S31}$$

where $\delta$ is small. In the limit $\delta \rightarrow 0$, we can take $\epsilon = \delta$. At the lowest order in $\epsilon$, we then have $\epsilon^2/\delta^2 = 1$ and

$$S'_0(z)^2 \approx g_0(z) - \lambda \tag{S32}$$

where prime indicates derivative with respect to z. The solution is given by

$$S_0(z) = \pm i \int_0^z (\sqrt{g_0(w) - \lambda})dw \tag{S33}$$

At the next order of $\epsilon$, we have

$$S_1(z) = -\frac{1}{4}\ln(\lambda - g_0(z)) \tag{S34}$$

Assuming $(g_0(z) - \lambda) > 0$, the only way to satisfy boundary conditions with this approximation, the solution $u(z)$ at this order in $\epsilon$ has cosine and sine terms. To satisfy the boundary condition at $z = 1$, the sine term must vanish, which places the following condition on its argument:

$$\boxed{\frac{1}{\epsilon}\int_0^1 \sqrt{g_0(z) - \lambda}dz = n\pi} \tag{S35}$$

with $n = 1, 2, 3....$ In principle, this equation can be solved numerically for $\lambda$ (for each value of $n$) to provide estimates of the eigenvalues.

In particular for $n = 1$, the principal eigenvalue will satisfy:

$$\int_0^1 \sqrt{g_0(z) - \lambda_1}dz = \epsilon\pi = \frac{\pi D}{L^2 g_{max}} \tag{S36}$$

In the constant $g$ case: $g_0(z) \equiv 1$ and $g_{max} = r$, the principal eigenvalue is simply given by $\lambda_1 = r - \frac{\pi^2 D}{L^2}$ and the persistence criterion $\lambda_1 > 0$ translates to a critical relationship between growth, diffusion and length of the domain, classically known for this type of reaction-diffusion models, as critical patch size:

$$\lambda_1 > 0 \iff L > \pi\sqrt{\frac{D}{r}}.$$

In general, for $g(z)$ spatially varying, we must compute this criterion according to expression S36.

#### S3. The discrete-space approach to computing $\lambda_1$

By considering a suitable discretization of the patch $[0, L]$, with $M + 2$ equally spaced points

$$z_i = h_z i, \quad i = 0, ..., M + 1$$

and

$$h_z = \frac{L}{M+1},$$

we can rewrite the eigenvalue equation

$$\frac{d^2 S}{dz^2} + \frac{g(z)}{D} S \quad = \quad \lambda S \tag{S37}$$

where $z \in [0, L]$ and $S(0) = 0$ and $S(L) = 0$, by using second-order centered finite differences to approximate the diffusion in each interior point of the discretized patch.

$$\frac{\partial^2 S_i}{\partial z} \approx \frac{u_{i-1} - 2S_i + S_{i+1}}{h_z^2} + O(h_z^2), \tag{S38}$$

with $S_i = S(h_z i)$. Note that we exclude the points $z_0$ and $z_{K+1}$ because the boundary conditions make the value of these points 0.

With this, we can write the discrete version of the eigenvalue equation 6 in each of the non-boundary points of our discretization as:

$$\begin{cases} \lambda_i S_i = \frac{g_i}{D} S_i + \frac{1}{h_z^2}(-2S_i + S_{i+1}), & \text{i=1} \\ \lambda_i S_i = \frac{g_i}{D} S_i + \frac{1}{h_z^2}(S_{i-1} - 2S_i + S_{i+1}), & \text{i=2,...,M-1} \\ \lambda_i S_i = \frac{g_i}{D} S_i + \frac{1}{h_z^2}(S_{i-1} - 2S_i), & \text{i=M} \end{cases} \tag{S39}$$

where $g_i = g(z_i)$ is the value of the growth rate of the mutant in the point $z_i$. We can write:

$$g(z_i) = g_{max} g_0(z_i)$$

***Rescaling of space***. Considering a rescaling of space, from $[0, L]$ to $[0, 1]$, we can consider the new system where now each discrete unit of space is given by $h = 1/(M + 1)$, so $1/h^2 = (M + 1)^2$, and the new $z_i = \frac{i}{M+1}, i = 1, ..M$.

$$\begin{cases} \lambda_i u_i = \frac{g_i L^2}{D} u_i + (M+1)^2(-2u_i + u_{i+1}), & \text{i=1} \\ \lambda_i u_i = \frac{g_i L^2}{D} u_i + (M+1)^2(u_{i-1} - 2u_i + u_{i+1}), & \text{i=2,...,M-1} \\ \lambda_i u_i = \frac{g_i L^2}{D} u_i + (M+1)^2(u_{i-1} - 2u_i), & \text{i=M} \end{cases} . \tag{S40}$$

Note, in this new spatial variable in $[0, 1]$, $g_i$ at each point $z_i$ corresponds to $g(z_i L)$ from the original function in $[0, L]$. Under the assumption:

$$g_i = g_{max} g_0(z_i L),$$

we can define

$$\epsilon = \frac{g_{max} L^2}{D},$$

and the above system can be rewritten in matrix form:

$$\lambda u_\lambda = (\Omega_0 + \epsilon G_0) u_\lambda, \tag{S41}$$

where $\Omega_0$ is the $M \times M$ square matrix for diffusion:

$$\Omega_0 = (M+1)^2 \begin{bmatrix} -2 & 1 & & & 0 \\ 1 & -2 & 1 & & \\ & \ddots & \ddots & \ddots & \\ & & 1 & -2 & 1 \\ 0 & & & 1 & -2 \end{bmatrix},$$

and $G_0$ is the $M$×M matrix for growth at each point in space.

$$G_0 = \frac{1}{g_{max}} \begin{bmatrix} g_1 & & & & 0 \\ & g_2 & & & \\ & & \ddots & & \\ & & & \ddots & \\ 0 & & & & g_M \end{bmatrix}$$

Our aim is to compute the eigenvalues of this system. For this we can resort to a perturbation approach. Assuming that $\epsilon \ll 1$, we expand our eigenvalue equation around the diffusion matrix:

$$(\Omega_0 + \epsilon G_0) |u_n\rangle_\epsilon = \lambda_n^\epsilon |u_n\rangle_\epsilon . \tag{S42}$$

Regarding the terms that appear in this equation, $\epsilon$ is a small positive real number, $0 < \epsilon << 1$,

$$|u\rangle_\epsilon = |u^0\rangle + \epsilon |u^1\rangle + \epsilon^2 |u^2\rangle + ...$$

is a Taylor series expansion of the normalized vector $|u\rangle$ and

$$\lambda_n^\epsilon = \lambda^0 + \epsilon\lambda^1 + \epsilon^2\lambda^2 + ...$$

a Taylor expansion of $\lambda_n$.

*S3.1. First order approximation*

If we expand equation S42, considering only the first-order terms (the ones with coefficient $\epsilon$ or 1), we obtain

$$(\Omega_0 + \epsilon G_0)(|u^0\rangle + \epsilon |u^1\rangle) = (\lambda_n^0 + \epsilon\lambda_n^1)(|u^0\rangle + \epsilon |u^1\rangle), \tag{S43}$$

and after some simplifications, we reach the equation

$$\Omega_0 |u^0\rangle + \epsilon\Omega_0 |u^1\rangle + \epsilon G_0 |u^0\rangle + \epsilon^2 G_0 |u^1\rangle = \lambda_n^0 |u^0\rangle + \epsilon\lambda_n^0 |u^1\rangle + \epsilon\lambda_n^1 |u^0\rangle + \epsilon^2\lambda_n^1 |u^1\rangle . \tag{S44}$$

To make a first-order approximation using perturbation theory, we must now equal the terms multiplied by the same powers of $\epsilon$ from both sides, ignoring the terms with powers of $\epsilon$ larger than one:

$$\begin{cases} \Omega_0 |u^0\rangle = \lambda_n^0 |u^0\rangle \\ \epsilon(\Omega_0 |u^1\rangle + G_0 |u^0\rangle) = \epsilon(\lambda_n^0 |u^1\rangle + \lambda_n^1 |u^0\rangle) \end{cases} . \tag{S45}$$

The first equation is always verified as it relates to the unperturbed system. Regarding the second equation, in order to simplify it, we can consider its inner product with $|u^0\rangle$, by multiplying on the left by $\langle u^0|$:

$$\begin{aligned} &\langle u^0| \Omega_0 |u^1\rangle + \langle u^0| G_0 |u^0\rangle = \langle u^0| \lambda_n^0 |u^1\rangle + \langle u^0| \lambda_n^1 |u^0\rangle \iff \\ &\iff \langle u^0| (\Omega_0 - \lambda_n^0) |u^1\rangle + \langle u^0| G_0 |u^0\rangle = \langle u^0| \lambda_n^1 |u^0\rangle \\ &\iff \lambda_n^1 = \langle u^0| G_0 |u^0\rangle , \end{aligned} \tag{S46}$$

which leads to the first-order term of the approximation of the eigenvalue $n$. The last step of this equation is done by noting that $\langle u^0| (\Omega_0 - \lambda_n^0) |u^1\rangle = 0$ since the system is unperturbed and $\langle u^0| \lambda_n^1 |u^0\rangle = \lambda_n^1 \langle u^0\rangle u^0 = \lambda_n^1$, due to the orthonormality of $\langle u^0|$ and $|u^0\rangle$ and noting that the eigenvalue is a number. Finally, to obtain the approximation of the $n^{th}$ largest eigenvalue, we do

$$\lambda_n = \lambda_n^0 + \epsilon\lambda_n^1 = \lambda_n^0 + \epsilon \langle u^0| G_0 |u^0\rangle . \tag{S47}$$

*Using the eigenvalues of the diffusion and growth matrix.* To obtain the values of $\lambda_n^0$ and $|u^0\rangle$, one can look at the properties of the $M \times M$ diffusion matrix $\Omega_0$, also known as a tridiagonal Toeplitz matrix (Deng 2021) and find that the eigenpairs $(e_j, u_j)$, with $u_j = (u_{j,1}, ..., u_{j,M})$ are

$$\begin{aligned} e_j &= (M+1)^2(-2 + 2\cos(j\pi h)) \\ u_{j,i} &= k\sin(j\pi i h), \qquad h = \frac{1}{M+1} \qquad\qquad j, i = 1, ..., M. \end{aligned} \tag{S48}$$

We can obtain the constant $c$ that appears in the following equations by

$$c = \sum_{i=1}^{M} \sin^2\left(\frac{ni\pi}{M+1}\right) = \frac{M+1}{2}, \tag{S49}$$

a constant that is used to normalize the eigenvectors so that we don't have different results for different discretizations of the spatial domain.

With these considerations, we obtain

$$\lambda_n^\epsilon = \lambda_n^0 + \frac{\epsilon}{g_{max}} \sum_{i=1}^{M} \frac{g_i}{c} \sin^2\left(\frac{ni\pi}{M+1}\right) = e_n + \frac{\epsilon}{g_{max}} \sum_{i=1}^{M} \frac{g_i}{c} \sin^2\left(\frac{ni\pi}{M+1}\right), \tag{S50}$$

where $c$ is the positive normalization constant that depends on our discretization (Eq. S49): The principal eigenvalue corresponds to $n = 1$:

$$\boxed{\lambda_1 \approx (M+1)^2\Big[-2 + 2\cos\left(\frac{\pi}{M+1}\right)\Big] + \frac{\epsilon}{c g_{max}} \sum_{i=1}^{M} g_i \sin^2\left(\frac{i\pi}{M+1}\right)}. \tag{S51}$$

Replacing $\epsilon = \frac{g_{max}L^2}{D}$ we obtain

$$\lambda_1 \approx (M+1)^2\Big[-2 + 2\cos\left(\frac{\pi}{M+1}\right)\Big] + \frac{L^2}{Dc} \sum_{i=1}^{M} g_i \sin^2\left(\frac{i\pi}{M+1}\right). \tag{S52}$$

Using a Taylor approximation of the cosine ($\cos(x) \approx 1 - \frac{x^2}{2!}$), which is reasonable for $M \to \infty$, we obtain:

$$\begin{aligned} \lambda_1 &\approx (M+1)^2\Big[-2 + 2(1 - \frac{\pi^2}{2(M+1)^2})\Big] + \frac{L^2}{Dc} \sum_{i=1}^{M} g_i \sin^2\left(\frac{i\pi}{M+1}\right) \\ &= -\pi^2 + \frac{L^2}{Dc} \sum_{i=1}^{M} g_i \sin^2\left(\frac{i\pi}{M+1}\right) \end{aligned} \tag{S53}$$

Now, replacing $c = \frac{M+1}{2}$ and reverting back to the original spatial scale (the function $g$ and *sin* now rescaled to $[0, L]$) we obtain:

$$\boxed{\lambda_1 = -\frac{\pi^2 D}{L^2} + \frac{2}{M+1} \sum_{i=1}^{M} g_i \sin^2\left(\frac{i\pi L}{M+1}\right).} \tag{S54}$$

Using the integral approximation

$$\sum_{i=1}^{M} g_i \sin^2\left(\frac{i\pi L}{M+1}\right)\frac{L}{M+1} \approx \int_0^L g(z) sin^2\left(\frac{\pi z}{L}\right)dz$$

or

$$\sum_{i=1}^{M} g_i \sin^2\left(\frac{i\pi L}{M+1}\right) \approx \int_0^L g(z) sin^2\left(\frac{\pi z}{L}\right)dz\frac{M+1}{L},$$

we get the match with the expression (Eq. S22 from the continuous-space approximation in Section S1:

$$\lambda_1 \approx= -\frac{\pi^2 D}{L^2} + \frac{2}{L}\int_0^L g(z)\sin^2\left(\frac{\pi z}{L}\right)dz$$

*S3.2. Second order approximation*

To obtain this second order approximation for $\lambda_1$, we follow a similar procedure as in equation S43, but now including terms with $\epsilon^2$:

$$(\Omega_0 + \epsilon G_0 + \epsilon^2 G_0)(|u^0\rangle + \epsilon\,|u^1\rangle + \epsilon^2\,|u^2\rangle) = (\lambda_u^0 + \epsilon\lambda_n^1 + \epsilon^2\lambda_n^2)(|u^0\rangle + \epsilon\,|u^1\rangle + \epsilon^2\,|u^2\rangle). \tag{S55}$$

After some simplifications and isolating the terms with $\epsilon^2$, we obtain

$$\lambda_n^2 = \langle u^0|\,G_0\,|u^1\rangle\,, \tag{S56}$$

where $|u^1\rangle$ is the first order correction to the wave function given by

$$|u^1\rangle = \sum_{k\neq n}\frac{\langle k^0|\,G_0\,|u^0\rangle}{\lambda_n^0 - \lambda_k^0}\,|k^0\rangle\,, \tag{S57}$$

where $k$ is used to identify the other eigenvalues and eigenvectors of the diffusion matrix. Using this, we can explicitly write the second order term of $\lambda_n^2$ as

$$\lambda_n^2 = \langle u^0|\,G_0\sum_{k\neq n}\frac{\langle k^0|\,G_0\,|u^0\rangle}{\lambda_n^0 - \lambda_k^0}\,|k^0\rangle = \sum_{k\neq n}\frac{|\langle k^0|\,G_0\,|u^0\rangle\,|^2}{\lambda_n^0 - \lambda_k^0}. \tag{S58}$$

In this case, the expression for $\lambda_1^2$, with the properties of the diffusion matrix, is

$$\lambda_1^2 = \frac{1}{c^2 g_{max}^2}\sum_{k=2}^{M}\sum_{i=1}^{M} g_i \sin\left(\frac{\pi i}{M+1}\right)\sin\left(\frac{\pi ik}{M+1}\right)\left(\frac{\sum_{i=1}^{M} g_i \sin\left(\frac{\pi i}{M+1}\right)\sin\left(\frac{\pi ik}{M+1}\right)}{2\cos\left(\frac{\pi}{M+1}\right) - 2\cos\left(\frac{k\pi}{M+1}\right)}\right), \tag{S59}$$

with $c^2$ coming from the same normalization constant as in equation S50.

The second-order approximation is then given by: $\lambda_1 = \lambda_1^0 + \epsilon\lambda_1^1 + \epsilon^2\lambda_1^2$ which in our system writes as:

$$\boxed{\lambda_1 \approx (M+1)^2\left[-2 + 2\cos\left(\frac{\pi}{M+1}\right)\right] + \frac{\epsilon}{c g_{max}}\sum_{i=1}^{M} g_i \sin^2\left(\frac{i\pi}{M+1}\right) + \epsilon^2\lambda_1^2}. \tag{S60}$$

To make the expression fully explicit we must substitute $\epsilon = \frac{g_{max}L^2}{D}$, revert to original space $[0, L]$, and simplify. It is noteworthy to remark that unlike the first-order approximation for $\lambda_1$, this second-order approximation provides a global measure of strain fitness, whose difference between two strains would involve both the strain growth functions and the common diffusion coefficient (a phenomenon observed in numerical simulations, where selection outcomes could flip with the diffusion coefficient for fixed growth functions). However, the weight of diffusion in equation S60 is usually very small, and, thus, in most cases, the predictions remain unchanged if we neglect this higher-order term.

#### S4. $\lambda_1$ as a measure for comparative fitness

##### *S4.1. Two general cases*

In general, we can use the expression for $\lambda_1$ to approximate the fitness of any strain in the system, and compare any two strains or multiple strains relative to each other. Below we illustrate this comparison based on the first-order approximation for $\lambda_1$ (Equation S22).

*Strains with same D.* If the diffusivity constant $D$ is the same, then the difference in fitness between any two strains becomes simply:

$$\lambda_1(i) - \lambda_1(j) = \frac{2}{L}\int_0^L \Big(g_i(z) - g_j(z)\Big)\sin^2\left(\frac{\pi z}{L}\right) dz. \tag{S61}$$

If the above difference is positive strain $i$ should be superior to $j$, and ultimately persist by excluding $j$ everywhere in space if furthermore its $\lambda_1(i) > 0$. If two strains have both negative $\lambda_1$, they will both go to extinction everywhere, but the one with higher $\lambda_1$ will go to extinction more slowly.

*Strains with different $D_i$.* In this case, strains would be allowed to vary both in motility traits and growth traits, and interesting selection outcome could result, for example if these traits are correlated, in positive or or negative (trade-off) manner. In general, the difference in fitness between two strains in this case is given by:

$$\lambda_1(i) - \lambda_1(j) = \frac{\pi^2}{L^2}(D_j - D_i) + \frac{2}{L}\int_0^L \Big(g_i(z) - g_j(z)\Big)\sin^2\left(\frac{\pi z}{L}\right) dz. \tag{S62}$$

Thus, under $D_i \neq D_j$, even with first-order $\lambda_1$ approximation, selection outcome between two strains becomes sensitive to diffusion parameters. The balance between growth and diffusion could be different between two strains, and coexistence may be more likely due to the contribution of diffusion asymmetries.

##### *S4.2. Mathematica codes and dynamic visualizations of the spatio-temporal dynamics of multiple mutants*

To illustrate the model dynamics, and the predictive value of $\lambda_1$, we include two Mathematica Notebooks

- Notebook 1: Solutions of the PDE system (related to Figure 2 of the paper)
- Notebook 2: $\lambda_1$ and predicting selection (related to Figure 6 of the paper)

and two separate supplementary movie files depicting the spatio-temporal dynamics of multiple random mutants ($\alpha_i$,$\beta_i$), under a given drug-interaction landscape and a joint concentration gradient:

- Movie S1: Selection of the “pink” mutant
- Movie S2: Selection of the “purple” mutant (when diffusion rate of the “pink” increases).

In the case of mutants with the same diffusion coefficient, the dynamics converge to the ”pink” strain winning, whereas in the case when diffusion coefficients vary, it is another strain that outcompetes all the others. One can verify and inspect these model scenarios also using and interacting with the Mathematica Notebook 2 above.

#### S5. Supplementary figures

##### *S5.1. More complicated Atlas for more complex growth rate function G.*

Here, we use more complex nonlinear growth functions dependent on the two drug concentrations $d_1$ and $d_2$:

$$G(d_1, d_2) = \frac{1}{1 + d_1^2}\left[1 + \left(\frac{d_2}{1 + \beta\frac{d_1}{1+d_1}}\right)^2\right]^{-1} \tag{S63}$$

and in particular, $\beta < 0$ for synergistic and $\beta > 0$ for antagonistic drug interaction.

This particular form of $G$ has been previously used to empirically describe a wide range of drug-drug landscapes (Wood et al., 2014), though for our purposes it should be viewed only as a simple parameterization of different 2d surfaces. The parameter $\beta$ modulates the shape of the constant growth contours, with $\beta < 0$ yielding synergistic and $\beta > 0$ antagonistic drug interactions.

The results are presented in Figure S2, which summarizes outcomes of selection on these four types of mutants, when they face different types of 2-drug concentration variation over space. We considered six different drug distributions over space to represent spatial heterogeneity:

1. Linearly-varying drug concentrations in opposite directions;
2. Linearly-varying drug concentrations in same direction;
3. Tangential U-shaped drug concentrations in opposite directions;
4. Intersecting U-shaped drug concentrations in opposite directions;
5. Hierarchical U-shaped drug concentrations with central maximum;
6. Hierarchical U-shaped drug concentrations with central minimum;

under two types of drug interaction (synergistic and antagonistic) and two diffusive scenarios (low and high diffusion). The initial distributions of each mutant were shaped as the function $\sin\left(\frac{\pi z}{L}\right)$, so that the homogeneous Dirichlet boundary conditions were respected. The spatial equilibrium plotted in Figure S2 is obtained numerically by considering the system at $t = 1000$. For all the simulations, initiated populations with 99% wild type mutants and the remaining 1% distributed equally among the three resistant mutants.

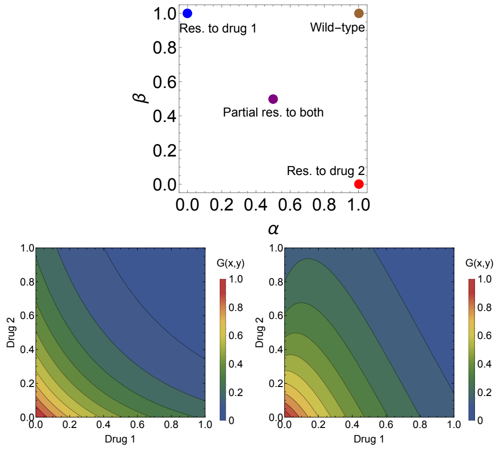


Figure S1: **The basis for the more complex atlas of multi-drug resistance evolution patterns over space. A.** The four canonical mutant types for resistance phenotypes to two drugs, distributed in the $(\alpha, \beta)$ space of rescaling parameters: blue - fully resistant to drug 1 and sensitive to drug 2; red - fully resistant to drug 2 and sensitive to drug 1; purple - intermediate resistance to both drugs; brown - wild-type, sensitive to both drugs. **B.** The 2 drug fitness landscapes used: synergistic (left) and antagonistic (right). For the synergistic drug interaction we assume $G^{syn}(d_1, d_2) = G(d_1, d_2)$ with $\beta = -1$, and the antagonistic drug interaction $G^{anta}(d_1, d_2) = G(d_1, d_2)$ with $\beta = 5$. These drug landscapes will be used to give rise to $g_i(z) = G(\alpha_i x(z), \beta_i y(z))$ as a function of two drug variation over space $x(z)$ and $y(z)$ with $z \in [0, 1]$. The relative fitnesses of the strains are hence dependent both on drug variation over space and on the details of the underlying growth landscape $G$.

We observe several final selection outcomes in these scenarios. For example cases when: i) the two single resistance mutants coexist (1a); ii) there is competitive exclusion and onle one single resistance mutant can persist everywhere over space (2a and 3a); iii) the intermediate multidrug resistance mutant is the final winner (1b, 2b); iv) Single- and multi-resistance coexist (5b). v) sometimes the outcome may depend on diffusion rate (3b). Coexistence seen here may be a transient phenomenon, although biologically relevant on realistic timescales. To strictly obtain pure coexistence we must make sure/verify the mean fitness is equal among coexisting variants. In the next figure (Fig.S3) we address this issue by exactly comparing fitness among mutants based on $\lambda_1$ when their growth rate varies over space.

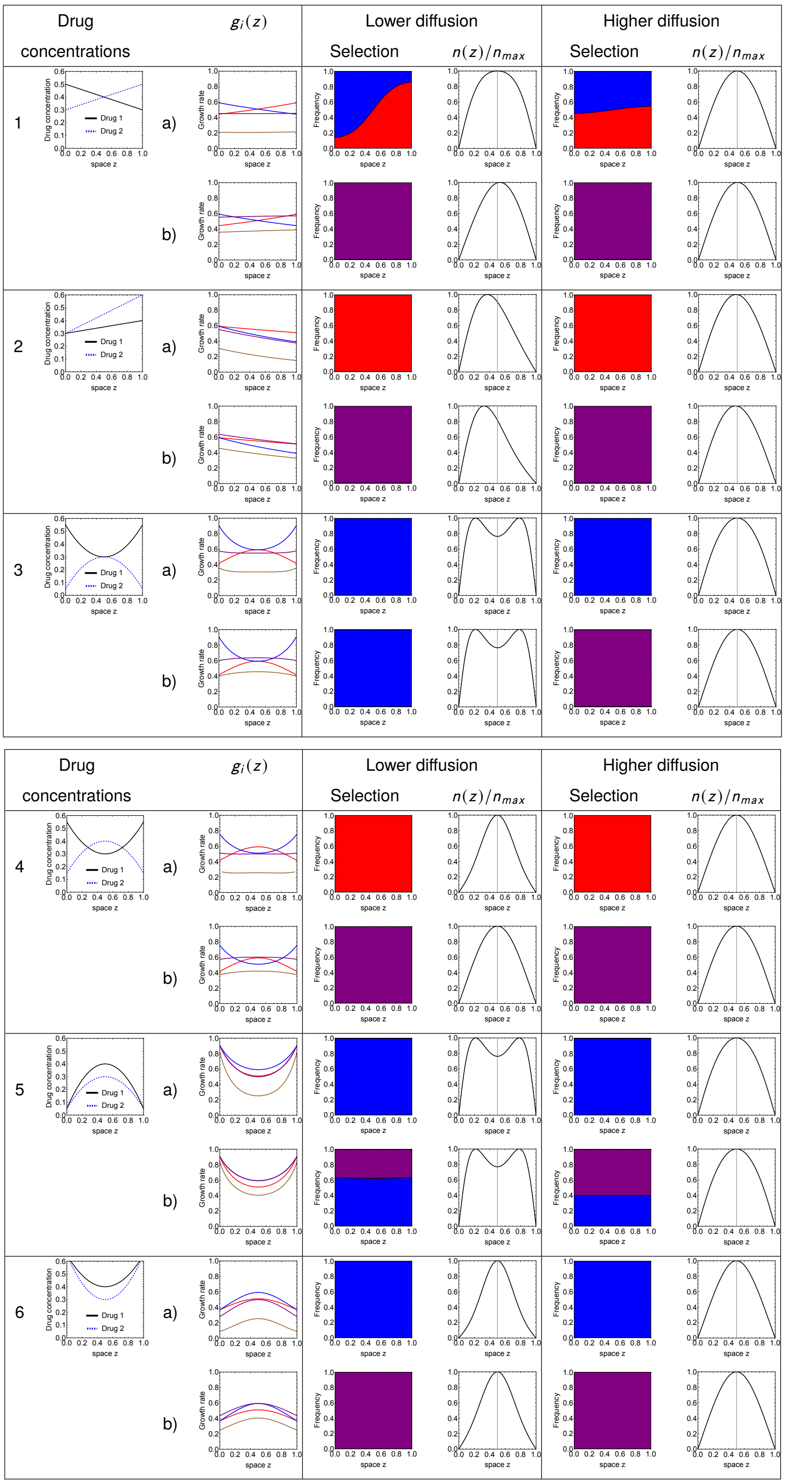


Figure S2: **An atlas for 2-drug resistance evolution in space under spatial heterogeneity**. We consider four available mutants each with different resistance phenotypes to two drugs, distributed in the $(\alpha, \beta)$ space: blue - mono-resistant to drug 1; red - mono-resistant to drug 2; purple - intermediate resistance to both drugs; brown - wild-type, sensitive to both drugs. Row a) of each scenario corresponds to a synergistic drug interaction ($G^{syn}$), and row b) to an antagonistic drug interaction ($G^{anta}$). Low diffusion corresponds to $D = 0.002$; High diffusion corresponds to $D = 0.02$. The $G$ landscapes were as in Figure S1.

| Scenario | $\lambda_1$ comparison | $\bar{g}$ comparison | Different prediction? |
|---|---|---|---|
| 1 a) | 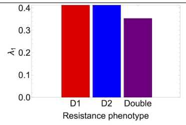 | 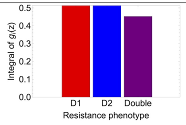 | |
| 1 b) | 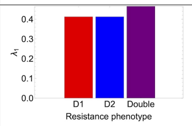 | 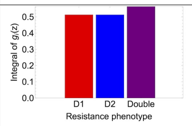 | |
| 2 a) | 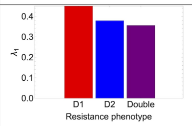 | 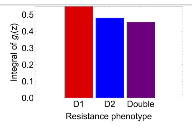 | |
| 2 b) | 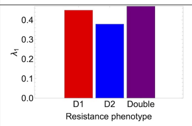 | 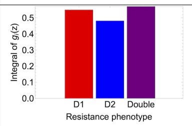 | |
| 3 a) | 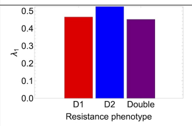 | 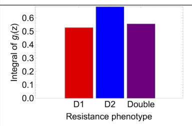 | |
| 3 b) | 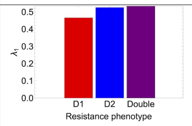 | 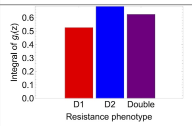 | Yes |
| 4 a) | 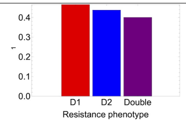 | 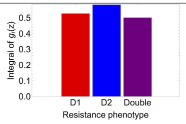 | Yes |
| 4 b) | 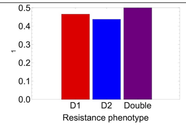 | 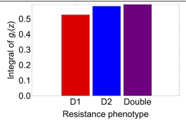 | |
| 5 a) | 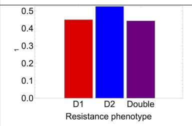 | 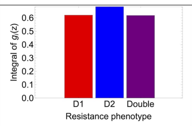 | |
| 5 b) | 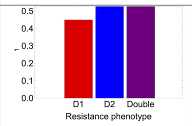 | 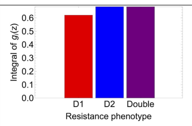 | |
| 6 a) | 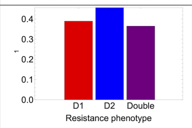 | 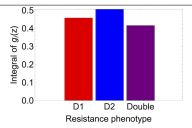 | |
| 6 b) | 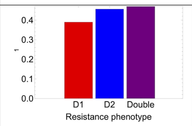 | 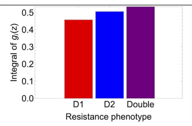 | |

Figure S3: **Comparing exact $\lambda_1$ predictions with numerical growth and diffusion simulations (Fig. S2) and $\bar{g}_i$ (spatially-averaged growth rate) rankings of mutants**. In most of these cases, the rankings obtained via $\lambda_1$ and mean $\bar{g}$ comparison predict the same selection outcome and match with what is observed in simulations.

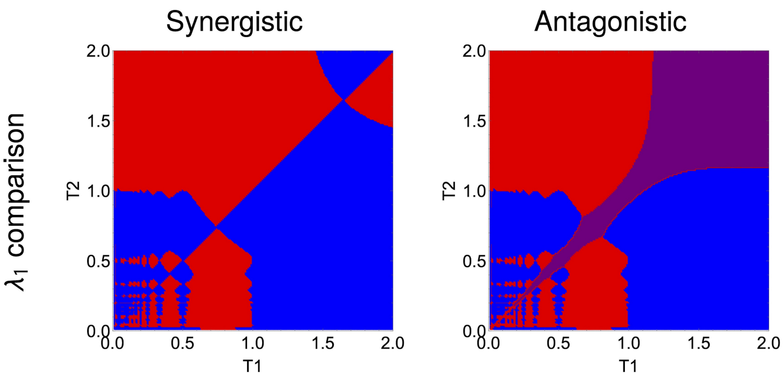


Figure S4: **The analogous selection outcomes under two periodic drugs for more complex drug interactions (Eq.S63)**. This figure shows the sensitivity of selection outcomes to the underlying growth functions, their nonlinearity, and their sensitivity and symmetry with respect to each drug. Compared to Figure 9 in the paper, the antagonistic drug combination here, for these two different drugs, does not enable as much the selection of the double-resistant mutant (purple region much smaller), still favouring mainly mono-resistance (blue or red strain wins), over all periodic multi-drug regimes considered.

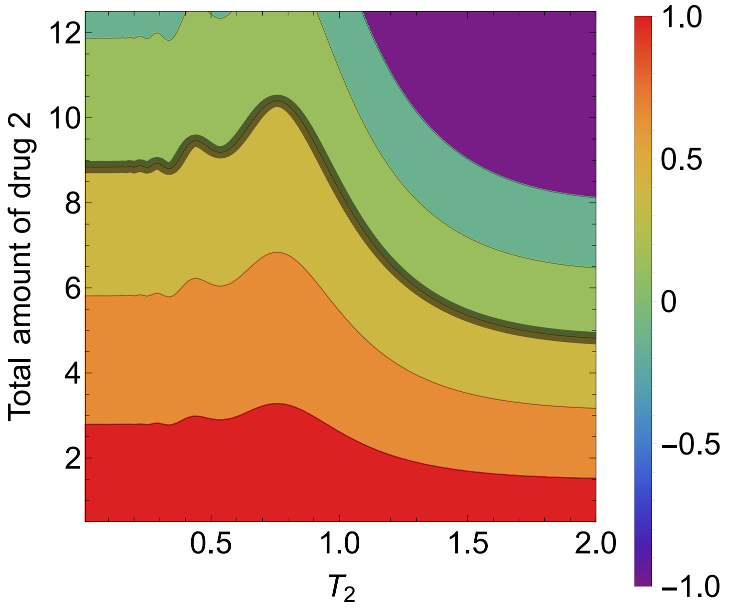


Figure S5: **Global fitness ($\lambda_1$) of the double resistance mutant as a function of drug-2 periodic variation.** We show a contour plot of the fitness of mutant $(0.5, 0.5)$, as a function of the total amount of drug 2 ($y(z)$) and the period of the drug distribution in space $z \in [0, 1]$. The periodic variation of drug 1 $x(z)$ was held fixed (including its total amount fixed at..), while drug 2 concentration $y(z)$ over space was varied by varying the period $T_2$. These parameters were fixed: $k_1 = A_1 = 0.5, T_1 = 0.8$, and the total amount of drug $y$ was rescaled. The assumed diffusion coefficient is $D = 0.02$. The drugs interact in a synergistic manner according to Eqs. 16 with $q = 0.5$. From this graph it is clear that there are different ways of drug-2 administration over space to effectively ensure elimination dynamics for the double resistance from the system, namely those combinations that make the $\lambda_1 < 0$. The contour-line separating regimes of possible growth from death is the $\lambda_1 = 0$ contour (highlighted in thicker gray line), potentially illustrating a minimal first trade-off curve for total amount vs. spatial period of drug 2 that will inhibit growth of the (0.5,0.5) variant.
